# An anaerobic Legionellales symbiont in *Anaeramoeba pumila*

**DOI:** 10.64898/2026.04.11.717937

**Authors:** Tengteng Zhang, Magdaléna Pavlátová, Shelby Williams, Dayana Salas-Leiva, Ivan Čepička, Andrew J. Roger, Jon Jerlström-Hultqvist

**Author notes:** Laboratory of Biodiversity and Evolution of Protozoa, College of Life Sciences, Shaanxi Normal University, Xi’an 710119, China (Tengteng Zhang). Department of Cell and Molecular Biology, BMC, Box 596, Uppsala Universitet, Sweden, SE-751 24 Uppsala, Sweden.

## Abstract

*Anaeramoeba pumila* is a free-living anaerobic amoeba and the smallest known member of the Anaeramoebae, a phylum characterized by elaborate membrane-bound symbiosomes housing sulfate-reducing bacterial symbionts. Here, we report a draft nuclear genome assembly of *A. pumila* LANTAAN and describe the discovery, genomic characterization, and metabolic reconstruction of *Candidatus* Centrionella anaeramoebae gen. nov., sp. nov., an obligate intracellular symbiont of *A. pumila* belonging to the order Legionellales. *Ca.* Centrionella is a rare anaerobic member of Legionellales, a lineage otherwise comprising aerobic intracellular pathogens. Its genome (1.52 Mbp, 1,249 genes) is highly reduced and encodes an entirely anaerobic metabolism centered on substrate-level phosphorylation, arginine fermentation, and hydrogen oxidation via a bidirectional [NiFe]-hydrogenase — metabolic strategies that parallel those independently evolved in the distantly related Anoxychlamydiales. The complete Dot/Icm type IVB secretion system is retained and likely mediates ongoing host manipulation, including via a large repertoire of predicted effector proteins. Strikingly, *Ca.* Centrionella has acquired eukaryotic Rac1-like GTPase genes from its host through horizontal gene transfer, with subsequent domain shuffling and duplication, that it may use to manipulate the cytoskeleton of its host. Unlike other *Anaeramoeba* symbionts, *Centrionella* localizes to the host microtubule-organizing center rather than a symbiosome, a localization consistent with cytoskeletal anchoring strategies described in other endosymbionts. The symbiosome, present in other *Anaeramoeba* species, appears to have been secondarily lost in *A. pumila*. A co-occurring *Desulfobacter* sp. LANTAAN, related to symbionts of other Anaeramoebidae, likely forms a tripartite syntrophic consortium by consuming hydrogenosomal fermentation end-products and supplying vitamin B_12_. Together, these findings illuminate the evolutionary transition in Legionellales from aerobic pathogenesis to anaerobic mutualism, providing a new model for the origins of intracellular symbiosis.

## Introduction

Amoebae interact with a multitude of different microbial organisms ranging from commensals to pathogenic organisms. In some instances, amoebae serve as reservoirs for organisms that cause infection in humans ^1^. In recent years, metagenomics and genome binning have led to the discovery of novel lineages that are presumed or are host-associated ^2–4^. However, many hosts and their associated microbes remain undescribed without cultivated members.

Novel host-associated bacterial lineages, such as Anoxychlamydiales ^5^, have adapted from aerobic ancestors to a lifestyle in oxygen-depleted environments such as marine sediments, yet their corresponding host lineages remain largely unknown. Likely, they colonize anaerobic protists that are yet to be cultivated or identified via single-cell sequencing efforts. Interestingly, the anoxychlamydial innovations parallel adaptations seen in anaerobic protists ^5^. Recent efforts have expanded the diversity and known host-range of another exclusively host-associated lineage, the Legionellales, via the description of multiple novel lineages that infect choanoflagellates, marine predatory protists related to symbionts that can act as pathogens in humans. Host-symbiont connections were established by fluorescence-activated cell sorting and single-cell sequencing of individual host cells with their symbionts ^4^. There are no similar high-throughput studies reported in anaerobic systems, and classical cultivation techniques remain the dominant route to new discoveries.

Anaeramoebids are a recently described phylum of free-living amoeboflagellates that inhabit oxygen-poor environments ^6^. Most *Anaeramoeba* species have an elaborate symbiosis-promoting membranous organelle, the symbiosome, that houses bacterial symbionts ^6^. Two different species of *Anaeramoeba* have been shown to house *Desulfobacter* sp., whose sulfate-reducing metabolism is a syntrophic match to the end-products produced by hydrogenosomes in *Anaeramoeba* ^7,8^. The symbiosome allows tight association and syntrophic interaction between the host hydrogenosomes and the symbionts, and still maintains surface connections that the symbionts likely require to access sulfate.

Interestingly, a novel anaeramoebid lineage of miniature cells, *Anaeramoeba pumila* isolate LANTAAN (hereafter *A. pumila*), forms a distinct phylogenetic lineage within *Anaeramoeba* ^9^. There are symbionts in *A. pumila,* and their phylogenetic origin is unclear, but in contrast to other Anaeramoebae, the symbiosome is not present, and symbionts are free in the cytoplasm in close proximity to the acentriolar microtubule-organizing centres (MTOCs) of the hosts. The symbiont cells are not closely associated with mitochondrion-related organelles (MROs) that appear to be randomly distributed in the cell ^9^. The metabolic capabilities of the MROs of *A. pumila* and how they relate to the symbionts remain unexplored.

Here we show that the symbiont of *A. pumila* is a gammaproteobacterium belonging to the Legionellales order UBA12402 (here described as *Candidatus* Centrionella anaeramoebae gen. nov., sp. nov., and referred to as *C. anaeramoebae* hereafter). We investigate the interaction between *A. pumila* and its symbiont using a combination of focused-ion-beam scanning electron microscopy (FIB-SEM) tomography, in situ localization techniques, genome sequencing, and comparative genomics of host and symbionts to shed light on their metabolic, infective, and evolutionary interplay with the host. We show that *Centrionella* has adapted to an anaerobic metabolism, unusual among Legionellales, and has potential syntrophic interactions with the host’s hydrogenosomal metabolism. *Centrionella* encodes the type IVb secretion system characteristic of other Legionellales and devotes a large proportion of its genome to large gene families some of which encode proteins with eukaryote-like domains. Among these, a Rac1-like GTPase of host origin has been duplicated within the *Centrionella* genome, yielding two copies with distinct domain architectures — potentially reflecting distinct strategies for manipulating host cell biology. Moreover, we show that a *Desulfobacter* sp. related to the symbionts of other anaeramoebids is a member of the *A. pumila*-associated microbiota and likely engages in a tripartite syntrophic interaction with *A. pumila* and *Centrionella*. These syntrophic interactions with *Centrionella* might be key to understanding the evolutionary route to the establishment and loss of the symbiosome in this *Anaeramoeba* lineage and offer clues to how symbiont replacement could occur.

## Results

### The symbionts are associated with the MTOC

Symbionts in *A. pumila* were observed at the center of the granuloplasm close to the MTOC and nucleus of the cell (Figure 1A). To clarify the relationship of the amoeba and its symbionts, we performed FIB-SEM of a single *A. pumila* cell. The cell was mononucleate and had four separate symbiont clusters comprised of rod-shaped cells (Figure 1B-F). Abundant mitochondrion-related organelles (>400) were globular in shape and did not cluster with the symbionts. As reported previously by Pavlátová and colleagues ^9^, no membranous symbiosome compartment was observed. Each of the symbiont clusters was associated with its own MTOC and arranged mostly on one side of the MTOC interface (Figure 1D-F). This is consistent with the co-localization of the nuclear and tubulin stain of the central position of the MTOC (Figure 1C). The four symbiont clusters had (8, 8, 7, 9) symbiont cells, respectively, and the latter two were closely associated with the nucleus.

**Figure 1:**
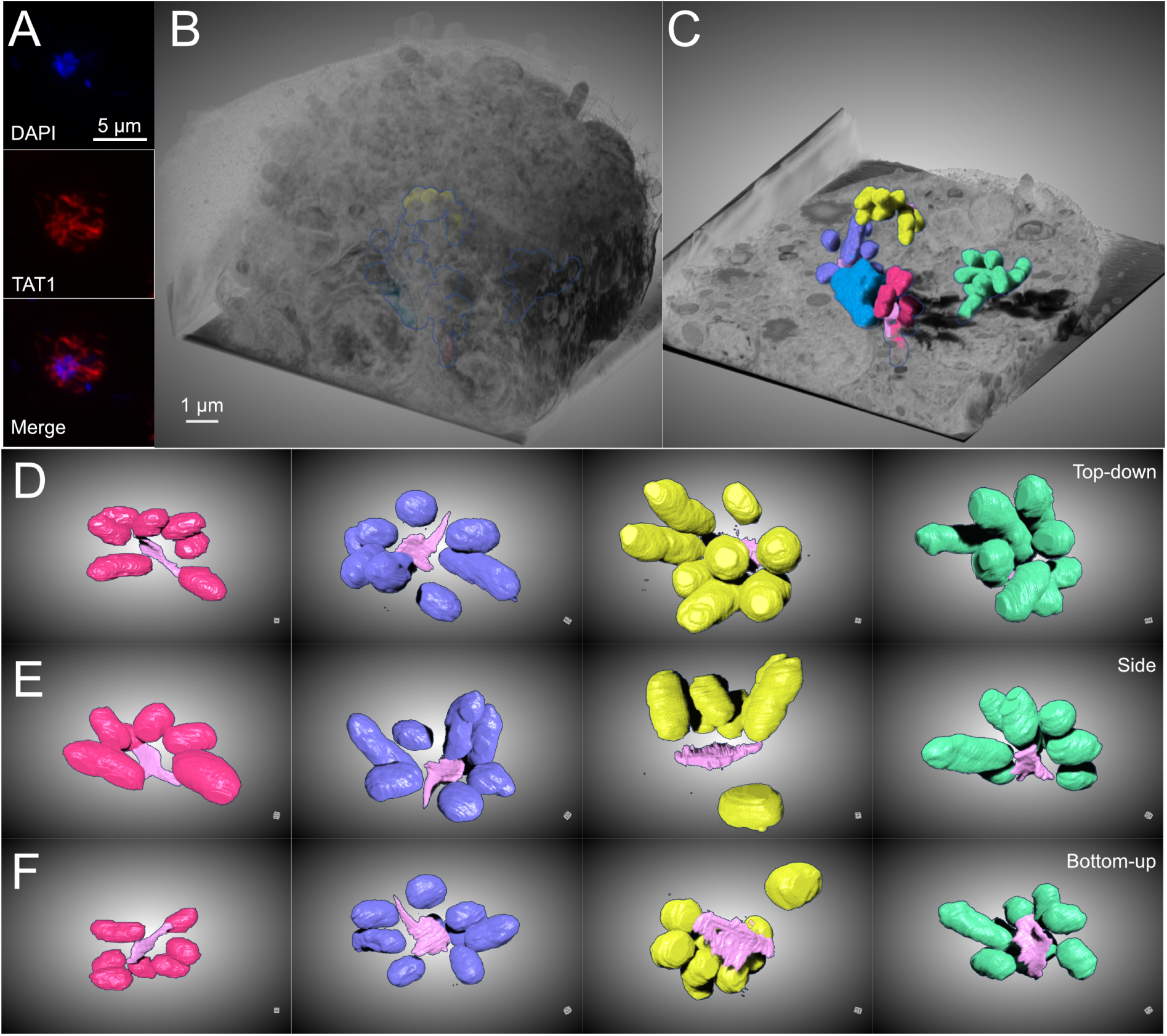
FIB-SEM of *A. pumila* cell with several symbiont clusters. **(A)** Immunofluorescence image of *A. pumila*. The blue signal indicates DNA (DAPI stain), and red signal indicates α-tubulin (TAT1) in the cells. Scale bar 5 µm. **(B)** Orthographic projection of *A. pumila* FIB-SEM volume. Symbiont clusters and the nucleus are shown in outline. Scale bar 1 µm. (**C)** Cut away of the cell showing the symbiont cluster and the nucleus. (**D)** Top-down, (**E)** side, and (**F)** bottom-up view of the four symbiont clusters. nucleus – blue, pink – microtubule organizing center, symbiont clusters – cerise, yellow, green, dark blue. The dataset and model are available via FigShare (10.6084/m9.figshare.31866142).

### The symbiont in *A. pumila* belongs to Legionellales

We used 16S rDNA profiling of the monoxenic *A. pumila* to investigate if any bacterial cells were consistently associated with the amoebae, as in Jerlström-Hultqvist et al (Figure 2A) ^8^. The planktonic bacteria showed were dominated by the genus *Vibrio* and *Draconibacterium*. The prokaryotic profile of the amoebae-enriched fractions was clearly distinct, and several groups, such as *Diplorickettsiaceae*, *Desulfobacter*, and *Carboxylicivirga,* not seen in the supernatants, were detected as prominent members. Interestingly, among the prominent members, the most amoeba-enriched fraction showed increased presence of *Diplorickettsiaceae,* whereas these were almost absent from the pellet fraction. In contrast, *Desulfobacter* and *Carboxylicivirga* are either depleted or unchanged, respectively. The cold-shocked enriched fraction was sequenced by nanopore sequencing, and the *A. pumila* metagenome assembly was manually binned into eukaryotic and prokaryotic bins using a combination of long-read, short-read, RNAseq, and BLAST search evidence. The metagenome assembly yielded high-coverage genomes of multiple bacterial genomes as well as the host genome (Table 1). We recovered circularized genomes of 11 prokaryotes, including a genome matching the sequence tag associated with *Diplorickettsiaceae,* a member of the order Legionellales of the phylum Pseudomonadota. Legionellales are all host- associated lineages, and most described species have been found to infect animals. Metagenomics has revealed a larger diversity of Legionellales lineages whose host affiliation is unclear. Recently, three lineages of Legionellales that were of unclear association were found to be associated with protists of the MAST-3 and choanoflagellate groups ^4,10^.

**Figure 2:**
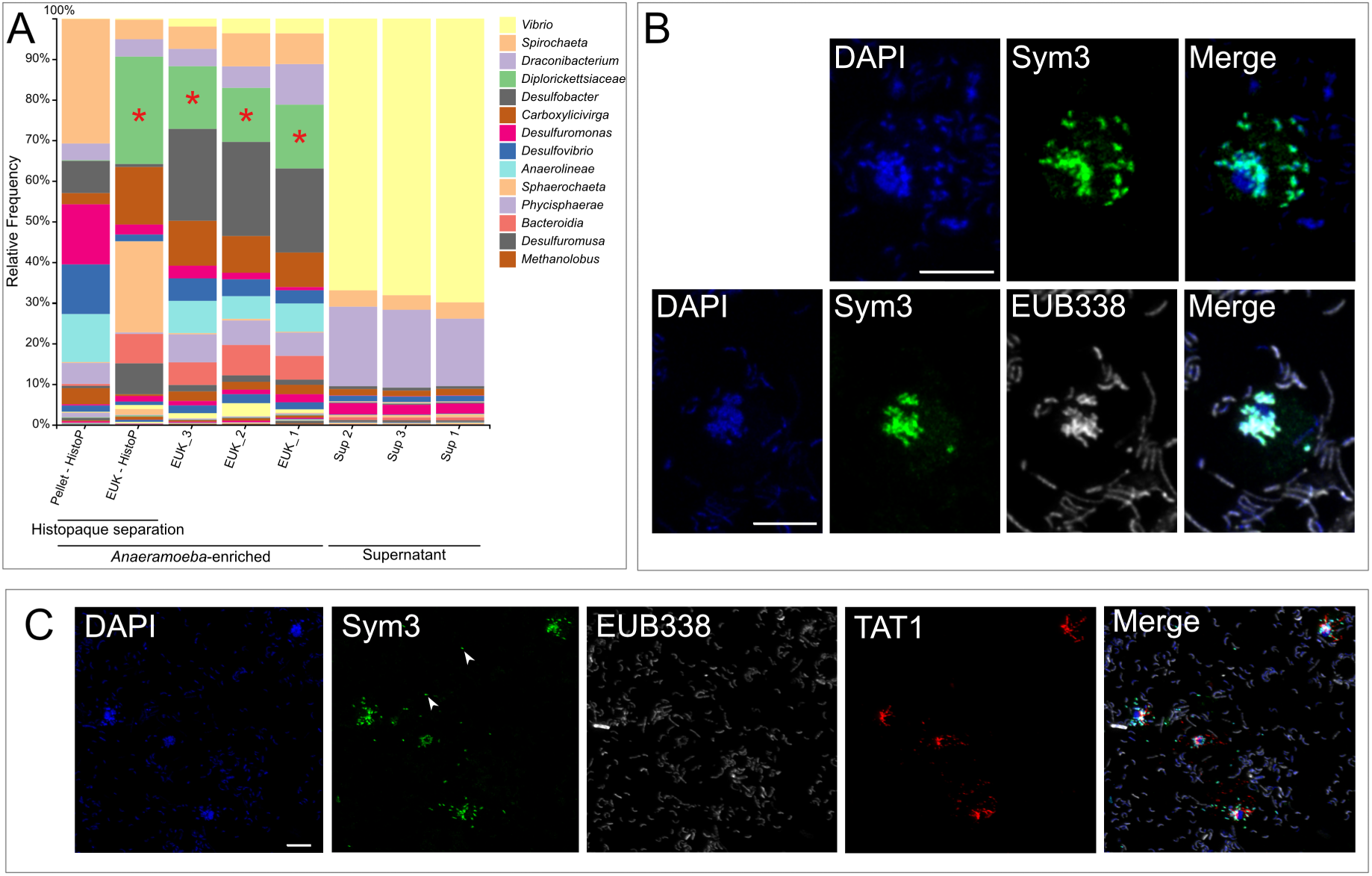
The endosymbiont bacteria in *A. pumila* belong to Legionellales. (**A)** 16S rDNA sequencing and analysis of *Anaeramoeba* enrichment and bacteria enrichment samples. Among them, EUK-Histop and Pellet-HistoP are samples enriched for eukaryotic and bacterial cells using a Histopaque discontinuous gradient, respectively. The other lanes are samples (EUK_X and Sup_X) where the cells and supernatant are separated by a cold shock procedure. The red asterisk (*) indicates endosymbiont bacteria. **(B)** Fluorescence in situ hybridization image of *A. pumila*. Blue, green, and white signals represent DAPI, Sym3 symbiotic bacteria probe, and universal bacterial probe, respectively. **(C)** Immunofluorescence and fluorescence in situ hybridization images of *A. pumila*. Blue, green, and white signals represent DAPI, Sym3 symbiotic bacteria probe (Sym3-Atto488), and universal bacterial probe (EUB338-Atto633) signals, respectively; red indicates the immunofluorescence signal of α-tubulin (TAT1 antibody detected by anti-mouse AlexaFluor 594); white arrows indicate potential symbiotic bacteria between cells. Scale bars 5 µm.

**Table 1:** Sequencing metrics and genome characteristics from the *A. pumila* genome compared to other sequenced *Anaeramoeba* genomes. # Data from the GCA_028826165.1 (*A. ignava*), GCA_027625915.1 (*A. flamelloides* Busselton2), and GCA_028892595.1 (*A. flamelloides* Schooner1) genome assemblies, as reported by ^8^. *Symbionts/amoeba reports the average nanopore coverage of the symbiont genome to coverage of the eukaryote assembly.

| Strain | *A. pumila*<br>LANTAAN | *A. ignava*<br>BMAN<sup>#</sup> | *A. flamelloides*<br>Busseton2<sup>#</sup> | *A. flamelloides*<br>Schooner1<sup>#</sup> |
| --- | --- | --- | --- | --- |
| Genome size (bp) | 25,818,118 | 42,843,398 | 254,083,758 | 259,028,678 |
| Coverage (Nano/Illumina) | 427 / 490 | 30 / - | 40 / - | 30 / - |
| # of scaffolds | 80 | 147 | 76 | 346 |
| N50 (bp) | 801,207 | 512,274 | 12,797,701 | 1,575,887 |
| L50 (n) | 10 |  |  |  |
| GC% | 52.74 | 19.24 | 23.63 | 24.75 |
| # of genes | 14,775 | 14,759 | 29,744 | 29,839 |
| # of tRNAs | 96 | 74 | 637 | 456 |
| Introns | 22 | 32,108 | 39,831 | 39,247 |
| Symbionts/amoeba* | 1.00 / 0.80 | 12.6 | 35.3 | 36.5 |

We designed a 16S rDNA-targeting probe Sym3 from the full 16S rDNA gene recovered from the genome, matching the sequence tag classified as *Diplorickettsiaceae*. Sym3 recognized bacterial cells visible via nuclear stain (Figure 2B). The average number of Sym3-positive symbionts per cell was 6 ± 3 (n=176). However, in some cultures, larger *A. pumila* amoebae with a higher number of symbionts were observed (>30 symbionts/cells). We noticed the Sym3 signal also outside of the cells (Figure 2C arrows). Whether this represents shed bacteria or material from cells that were damaged during microscopy sample preparation is unclear. Symbionts were often found close to the nucleus and the MTOC, as seen in the FIB-SEM analyses (Figure 2C).

### The symbionts of *A. pumila* belong to clade UBA12402

The full-length circularized genome of the symbiont is 1.52 Mbp, and codes for 1,249 genes (1,198 protein-coding) (Figure 3, Table 2). The coding density is 88.9% with 18 pseudogenes annotated. The genome shows evidence of recent activity by insertion sequences (IS) and other mobile elements, with a total of seventy intact or fragments of IS elements detected.

**Figure 3:**
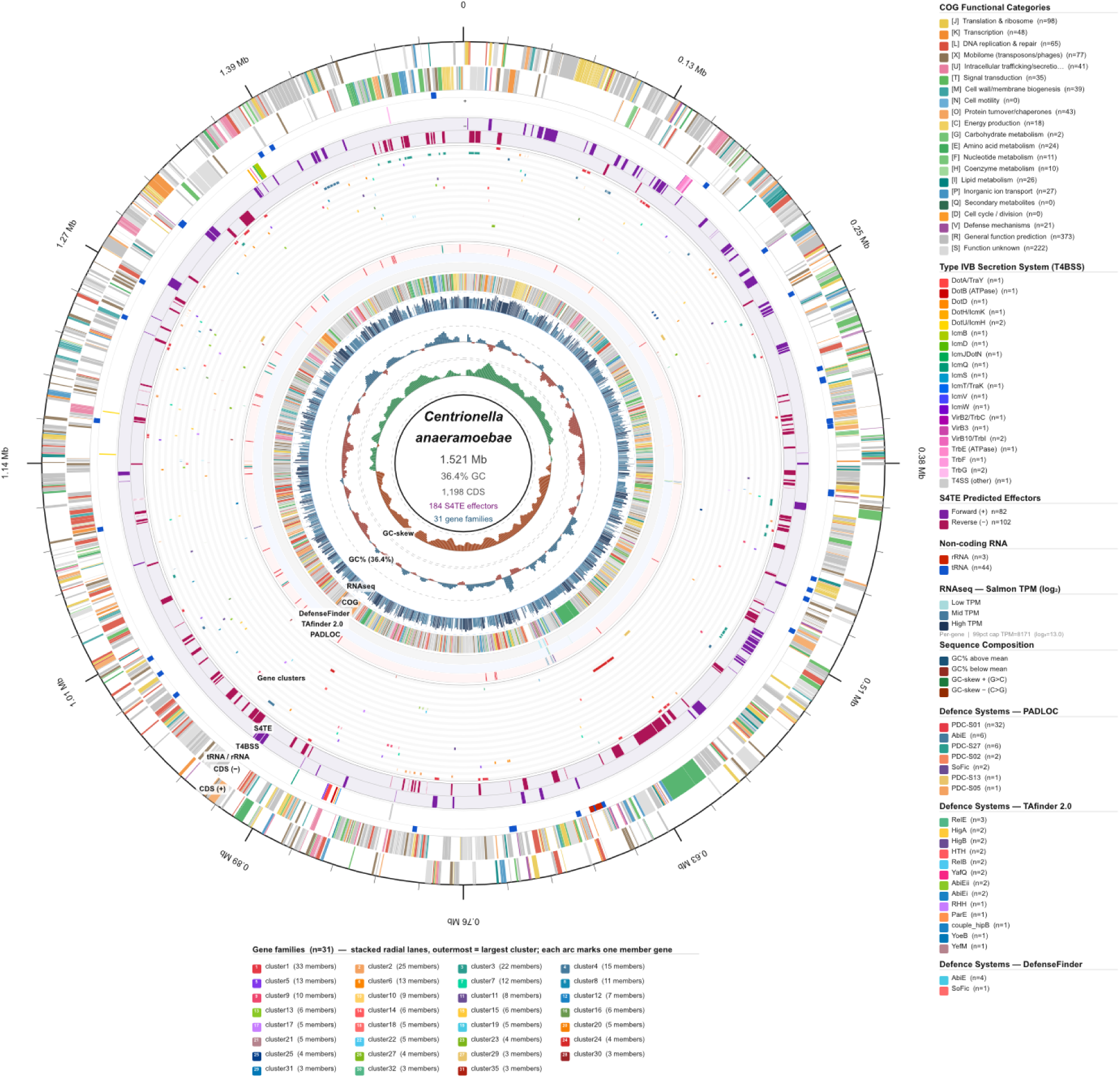
Genome atlas of *Centrionella anaeramoebae.* Circular genome map of the *C. anaeramoebae* chromosome (1,521,317 bp; 36.42% GC). Fourteen concentric tracks are arranged from outer to inner and are described below. All tracks are drawn at exact genomic coordinates; no positional smoothing or gene-length normalization was applied. Annotation comprises 1,198 protein-coding sequences, 44 tRNA genes, and 3 rRNA genes (16S, 23S, and 5S). **Track 1. Scale and tick marks.** The outermost ring shows the chromosomal coordinate scale. Major tick marks are placed every 500 kb with Mb labels; minor ticks are placed every 100 kb. Coordinates are numbered clockwise from the origin (position 0) and correspond directly to the GenBank accession coordinates. **Track 2 and 3. Protein-coding sequences**, forward (+) strand and, reverse CDS) (−) strand, respectively. **Track 4. Non-coding RNA genes**. This narrow ring marks the positions of the 44 tRNA genes (blue, #1155CC) and 3 rRNA genes (dark red, #CC2200). **Track 5. Type IVB secretion system (T4BSS) structural components**. Arcs represent the 22 T4BSS structural genes identified by pattern-matching against GenBank /product annotations. The Dot/Icm/VirB/TrbC/TrbE/TrbF/TrbG system is the *Legionella*-type conjugal transfer apparatus that mediates effector translocation. Each of the 20 component classes is assigned a distinct color. **Track 6. Predicted Type IV-secreted effectors (T4SE).** This track displays the 184 predicted effectors from S4TE. **Track 7. Predicted gene clusters.** Thirty-one predicted gene clusters (263 member genes in total) are rendered as stacked concentric lanes. Each cluster occupies a dedicated radial lane ordered from outermost (largest cluster) to innermost (smallest cluster). Within each lane, member genes are drawn as arcs at their exact genomic coordinates; white 0.4-point hairline strokes separate adjacent members. Each cluster is assigned a distinct color from a 31-member palette. **Track 8. Defense systems — PADLOC.** The first of three defense-system sub-rows (the outermost of the three) shows the 50 genes comprising seven defense systems predicted by PADLOC v.2.0.0^1^. Each system is assigned a distinct color. **Track 9. Defense systems — TAfinder 2.0.** The middle defense sub-row shows the 22 genes (11 toxin–antitoxin pairs) predicted by TAfinder 2.0^2^. The dataset contains 18 Type II and 4 Type IV TA systems. Each of the 13 TA families is assigned a distinct color; toxin and antitoxin components of each pair share the same family color. **Track 10. Defense systems — DefenseFinder.** The innermost defense sub-row shows the 5 genes (comprising 3 systems) predicted by DefenseFinder^3^ : AbiE_1 (2 genes), AbiE_2 (2 genes), and SoFic_3 (1 gene). AbiE genes are colored sky blue (#48CAE4); SoFic is colored coral red (#FF6B6B). Four of these five genes (the AbiE loci) are also detected by PADLOC (**Track 8**), and two are co-identified by all three defense-prediction tools (PADLOC, TAfinder, and DefenseFinder). In total, the three tools collectively annotate 68 unique defense-associated loci across the genome (50 PADLOC + 22 TAfinder + 5 DefenseFinder, with overlaps removed). **Track 11. COG functional annotation ring.** A narrow ring repeats the COG color code for all 1,180 CDS (both strands combined), providing a rapid overview of functional composition around the chromosome. Colours are identical to those used in **Tracks 2** and **3**. COG categories were assigned by keyword matching against GenBank /product annotations; genes whose products could not be assigned to an information-processing, metabolic, or cellular-process category were classified as R (general function prediction only) or S (function unknown). **Track 12. Transcriptomic expression — Salmon TPM.** This histogram track displays per-gene transcript abundance estimated by Salmon quantification^4^. Values are expressed as transcripts per million (TPM) and are plotted on a log₂(TPM + 1) scale. To prevent extreme outliers from compressing the color gradient, values are capped at the 99th-percentile TPM (8,171; log₂ = 13.00), so that all expressed genes remain visually distinguishable. Of the 1,165 genes quantified, 1,050 had TPM > 0 (expressed) and 115 had TPM = 0 (not detected in this condition). Bar height encodes the normalized expression level (0–1 scale) and bar color transitions continuously from pale teal (#A8DADC) through steel blue (#457B9D) to deep navy (#1D3557) as expression increases. **Track 13. GC content.** GC percentage was calculated in 15-kb sliding windows with a 3-kb step (503 windows total), producing a mean GC of 36.41% (range 33.5–40.4%). The histogram baseline is set at the genome-wide mean; windows above the mean are drawn in dark blue (#1A5276) and windows below the mean are drawn in dark red (#922B21). **Track 14. GC-skew.** GC-skew, defined as (G − C) / (G + C), was calculated over the same 15-kb/3-kb windows as **Track 13** (range −0.300 to +0.378). The baseline is set at zero; windows with G surplus (positive skew) are drawn in dark green (#1A7A3C) and windows with C surplus (negative skew) are drawn in dark orange (#922B00). GC-skew transitions correspond to the putative replication terminus (negative-to-positive transition) and origin of replication (positive-to-negative transition).

**Table 2:** Genome characteristics of the *Centrionella anaeramoebae*. Data is derived from NZ_CP092900.1 (*Candidatus* Comchoanobacter bicostacola), GCA_000815025 (*Coxiella* endosymbiont of *Amblyomma americanum*), and GCA_900452735.1 (*Legionella pneumophila* strain Philadelphia-1 isolate AE).

| Strain | *Candidatus*<br>*Centrionella*<br>*anaeramoebae* | *Candidatus*<br>*Comchoanobacter*<br>*bicostacola* | *Coxiella*-like<br>endosymbiont<br>(CLEEA) | *Legionella*<br>*pneumophila*<br>strain<br>Philadelphia-1 |
| --- | --- | --- | --- | --- |
| Genome size (bp) | 1,521,317 | 1,007,335 | 656,801 | 3,409,143 |
| Coverage<br>(Nano/Illumina) | 431 / 609 | - / 8000 |  | 106x PacBio |
| # of scaffolds | circular | circular | Circular | circular |
| GC% | 36.4 | 39.5 | 34.6 | 38.5 |
| # of genes | 1,249 | 985 | 582 | 3,057 |
| # of RNAs | 51 | 42 | 45 | 79 |

We performed a phylogenomic analysis of the symbiont genome within the context of the Legionellales using the dataset from Hugoson *et al.* ^11^. The phylogeny of 109 proteins from 63 Legionellales genomes recovered the *A. pumila* symbiont in the larger group defined as “Aquicella” (Figure 4). GTDB-tk classified the genome as belonging to the provisional UBA12402 order and family with the same name in the genus JAFGHQ01, which mainly includes members with metagenome-assembled genomes. The only member of this clade with a host association is *Candidatus* Cochliophilus cryoturris, which is described from the testate amoeba *Cochliopodium minus* 9B ^12^. The 16S rDNA sequence of Ca. C. cryoturris shows 97% similarity to the UBA12402 genome GCA_018812595, which belongs to the UBA12402 family but to a different genus ^4,12^. Our phylogenomic analyses support the GTDB-tk results, showing that the symbiont clusters in a clade with annotated UBA12402 members. We describe the *A. pumila* symbiont as Candidatus Centrionella anaeramoebae sp. nov. in line with its association with the MTOC in *A. pumila* cells. The deep divergence of this clade prompts us to describe the provisional order UBA12402 as *Candidatus* Centrionellales n. ord, and provisional family UBA12402 as *Candidatus* Centrionellaceae n. fam and the JAFGHQ01 genus as *Candidatus* Centrionella n. gen, respectively.

**Figure 4:**
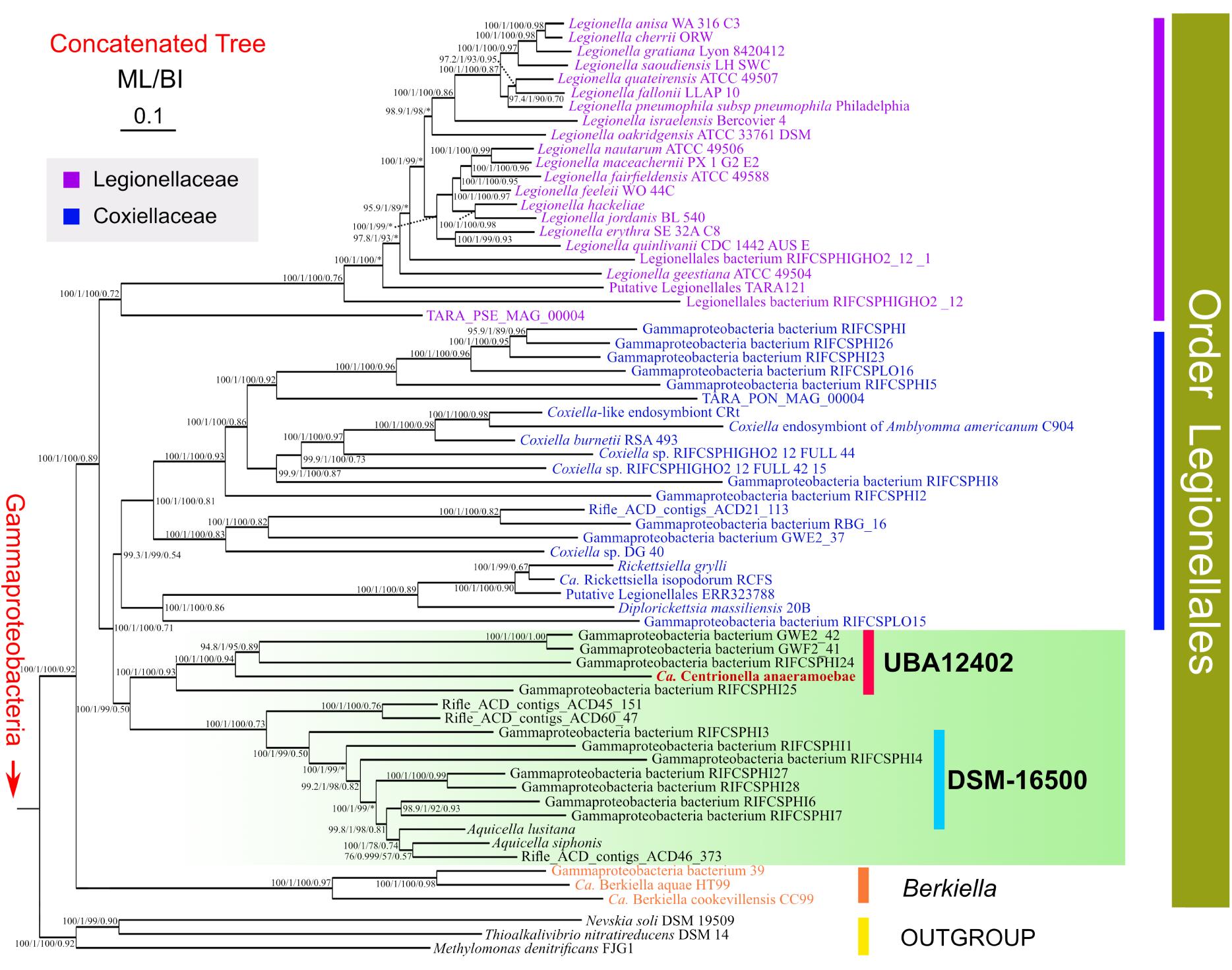
Phylogenomic analysis focusing on Gammaproteobacteria, inferred from 109 orthologous protein sequences with 29,139 amino acid sites. The maximum likelihood (ML) tree was constructed using IQ-TREE with the LG+C60+F+G4 model. The Bayesian inference (BI) tree was generated using PhyloBayes with the CAT+GTR model. Values at the branch nodes represent SH-aLRT, aBayes, standard confidence values, and BI confidence values, respectively. The asterisks “*” indicate inconsistent topologies between the ML and BI analyses. The order Legionellales, the families *Legionellaceae* and *Coxiellaceae. Berkiella, Aquicella* (green shading), UBA12402, DSM-16500 indicate important groups. The scale bar indicates a substitution rate of 10%.

### *Anaeramoeba pumila* has secondarily lost its symbiosome

The eukaryotic part of the assembly totalled 25.8 Mbp in length and was covered 303x on average (Table 1). We annotated 14,775 protein-coding genes and 206 non-coding RNA genes (110 rRNA loci, 96 tRNA), revealing a much higher coding density than in other *Anaeramoeba* genomes. This included 2 selenocysteine tRNAs, indicating that *A. pumila* can make selenoproteins. Remarkably, we only detected 22 introns in 19 genes across the genome compared to more than 30,000 introns per genome in other sequenced *Anaeramoeba* isolates (Table 1). This adds *A. pumila* as yet another eukaryotic lineage that has suffered massive intron loss ^13^. The splice sites in *A. pumila* were highly conserved at the 5’ end (GTTAGT) and at the branch point (CT(G/c)**A**C), as commonly seen in genomes that have lost most of their introns ^14–16^, whereas the 3’ splice site did not show any strong conservation apart from the standard AG (Figure S1). The overall GC-content in *A. pumila* (52.74%) is very different from that of other sequenced Anaeramoebae, which, so far, all have AT-rich genomes (Table 1).

We investigated the phylogenetic position of *A. pumila* using a 60-taxon phylogenomic dataset of 252 proteins that covers the diversity across Metamonada and most Eukaryotic supergroups (Obazoa, Amoebozoa, CRuMs, Malawimonadida, Discoba, Archaeplastida, Rhizaria Stramenopila, Alveolata) (Figure S2). The phylogeny recovers Anaeramoebae as a monophyletic group that branches sister to Parabasalia. *A. pumila* branches between the *A. ignava* and *A. flamelloides* groups. A more focused phylogenomic analysis of 251 proteins from 32 species, focusing on the Metamonada clade with Discoba as the outgroup, confirmed the results of the larger phylogenomic dataset. (Figure 5). *A. pumila* branches between *A. flamelloides* and *A. ignava*, with full support, and is most closely related to the former. In light of the phylogenomic results, several characteristics seen in other Anaeramoebae and missing in *A. pumila*, such as the lack of a symbiosome, an AT-rich genome, and a paucity of introns, appear likely to be secondary losses.

**Figure 5.**
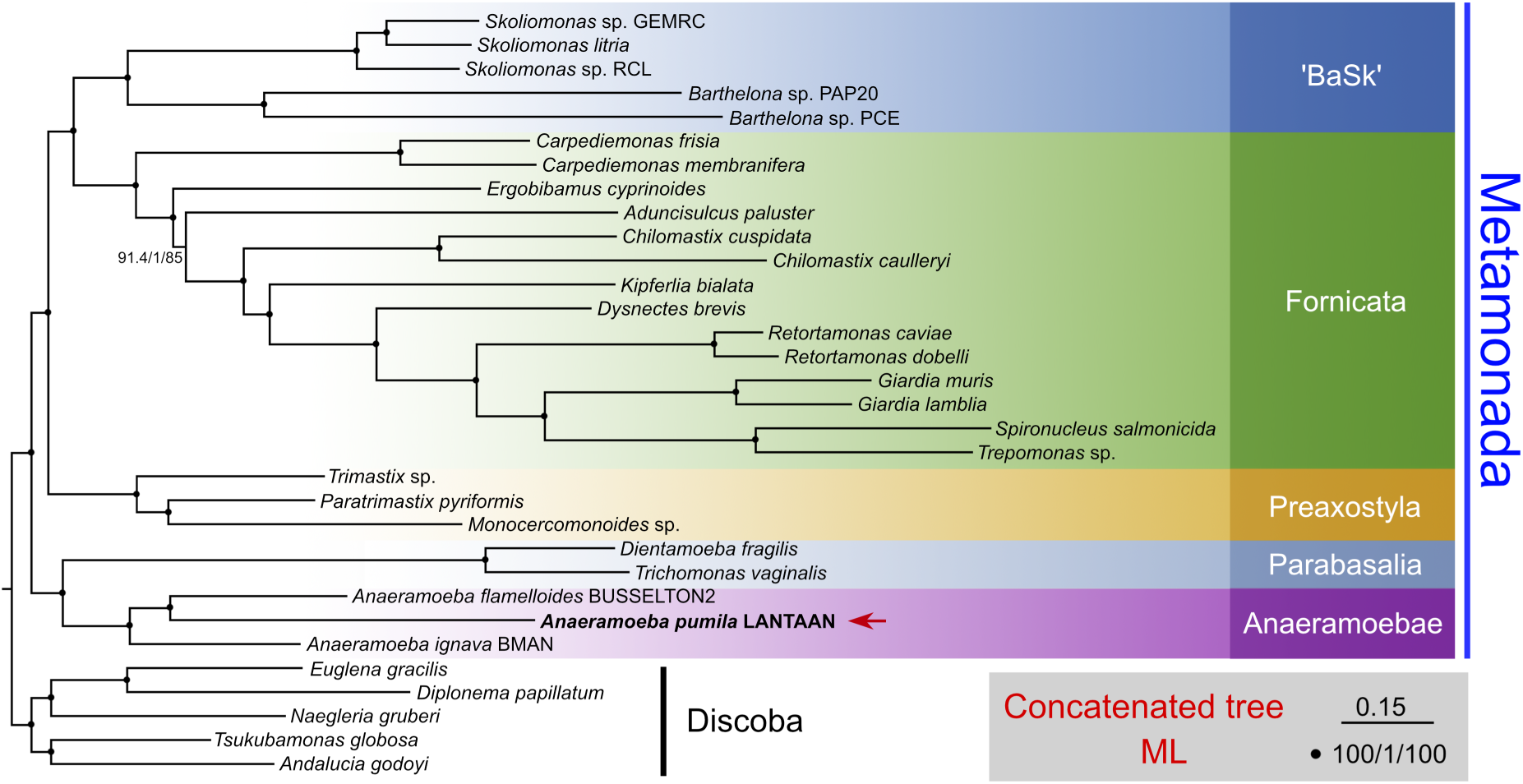
The maximum likelihood tree (ML) focusing on the *Anaeramoebae* was generated from 251 protein sequences with 69,041 amino acid sites. The ML tree was constructed using IQ-TREE with the LG+C60+F+G4 model. Values at the nodes represent SH-aLRT, aBayes, and standard confidence values, respectively. Solid dots indicate full confidence values. The scale bar indicates a substitution rate of 20%.

### *Anaeramoeba pumila* has hydrogenosomes

All Anaeramoebae investigated by electron-microscopy, except for *A. pumila*, house their symbionts in membrane vesicles that form symbiosomes. These structures are contiguous membrane structures with connections to the extracellular space and are intimately associated with hydrogenosomes. The symbioses in Anaeramoebae are thought to be sustained by the end-products of hydrogenosome metabolism — hydrogen, acetate, and propionate — the pathways for which were previously reconstructed in *A. ignava* and *A. flamelloides* ^7^. *A. pumila* lacks any traces of a symbiosome, and its symbionts show no clear association with its MROs ^9^. The presence of MROs was detected by immunofluorescence microscopy using a heterologous antibody raised against the mitochondrial Cpn60 from *Neocallimastix* (Figure S3). The antibody signal revealed tens to hundreds of point-like organelles depending on cell size, and did not show any apparent evidence of specific association with symbionts, MTOC, microtubules, or the nucleus (Figure S3), in keeping with the previous TEM observations by Pavlátová and colleagues ^9^. To clarify the predicted capacity of the MRO in *A. pumila,* we searched for MRO components using *Anaeramoeba* orthologs as well as unbiased methods (Figure 6). The reconstructed proteome and metabolism of *A. pumila* MROs show a similar predicted capacity to those of other Anaeramoebae ^7^. We detected core parts of the protein import and folding machinery, including the outer membrane protein Sam50, the intermembrane space Tim10 chaperone, and inner membrane proteins Pam16/18 and Tim17. We were unable to identify homologs of some import components identified in *A. flamelloides* and *A. ignava*, such as Tom22, Tom40, and Tim44. Crucial components of the protein folding chaperones, Hsp70 and Cpn60/10, as well as both components of the mitochondrial processing peptidase (MPP), were identified (Figure 6).

**Figure 6.**
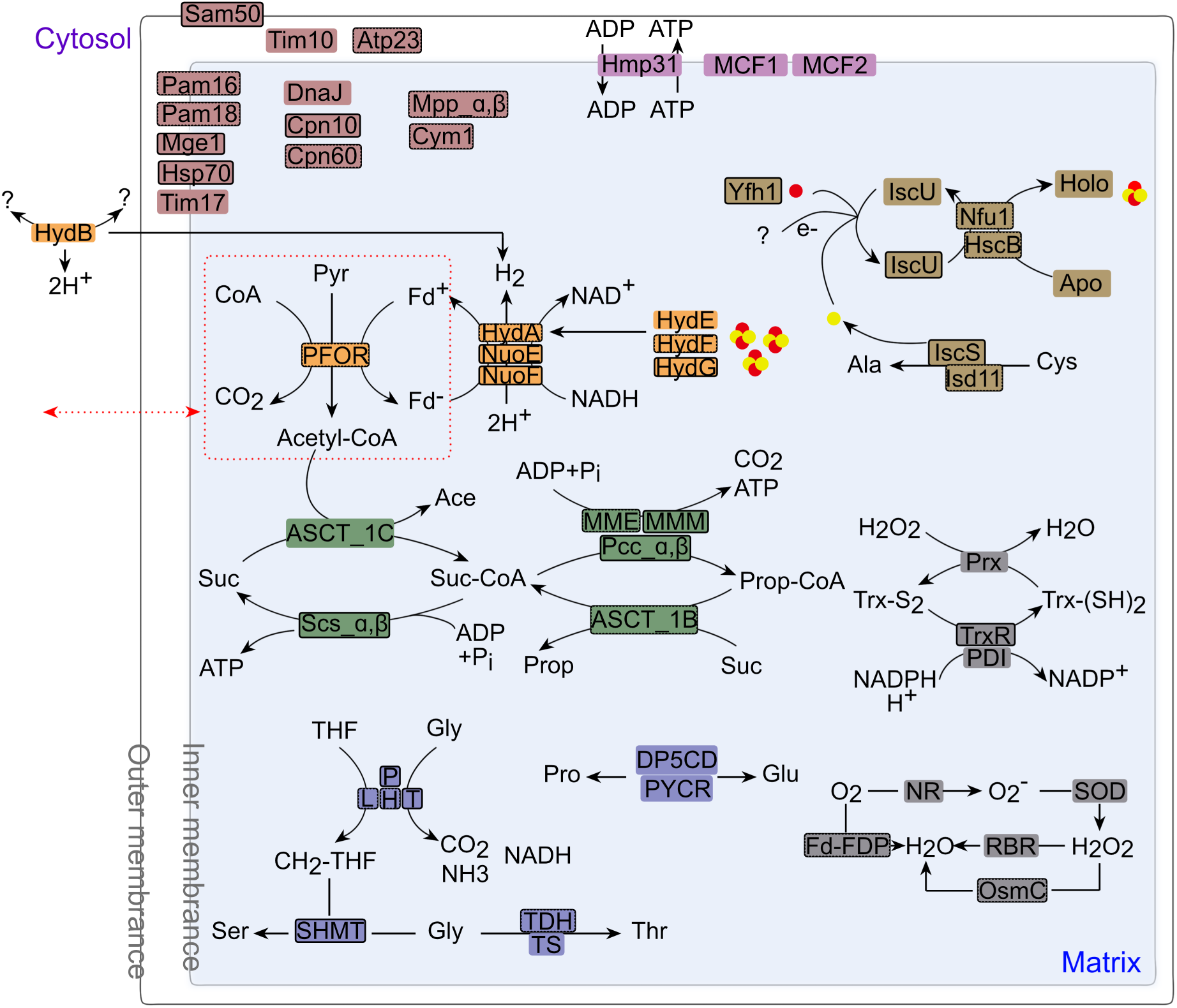
Metabolism prediction of mitochondrion-related organelles (MROs) in *Anaeramoeba pumila*. Proteins predicted to have MTS by all three software (Mitoprot, Mitofates, and TargetP) are outlined with solid borders. Proteins predicted to have MTS by only one or two software tools are outlined with dashed borders. Proteins with no MTS predicted by any of the three tools have no border. Unknown proteins or substrates are indicated by question marks. Solid yellow and red circles represent sulfur and iron, respectively. The proteins related to protein import and folding are highlighted in pink; mitochondrial carrier proteins are in light purple; proteins related to pyruvate and hydrogen metabolism are in orange; proteins involved in Fe-S cluster synthesis are in brown; proteins related to energy metabolism are in green; proteins related to amino acid metabolism are in blue; and proteins related to oxidative stress response metabolism are in gray. Pyr: pyruvate; CoA: coenzyme A; Acetyl-CoA: acetyl-coenzyme A; Ala: alanine; Cys: cysteine; Apo: Fe-S cluster-free protein; Holo: Fe-S cluster-containing protein; Ace: acetate; Suc: succinate; Suc- CoA: succinyl-CoA; Prop: propionate; Prop-CoA: propionyl-CoA; THF: tetrahydrofolate; Gly: glycine; CH2-THF: methylenetetrahydrofolate; Ser: serine; Thr: threonine; Pro: proline; Glu: glutamate; Trx-S2: oxidized thioredoxin; Trx-(SH)2: reduced thioredoxin. The red dashed box and red dashed arrows indicate that the reactions within the red dashed box also occur in the cytoplasm.

Fe-S cluster biogenesis is a central function of most studied MROs. The canonical mitochondrial system, the ISC-system, is present in *A. pumila,* and several components are predicted to localize to the MROs. Anaeramoebids are predicted to oxidize pyruvate via pyruvate:ferredoxin oxidoreductase (PFOR) to generate acetyl-CoA, CO2, and reduced ferredoxin (Fd^−^). Fd^−^ and NADH are likely transferred to an electron-confurcating multisubunit [FeFe]-hydrogenase that yields H2 as an end-product. The maturase proteins (HydEFG), which are needed to activate the [FeFe] hydrogenase, were all detected in *A. pumila*. The energy-generating pathways of *A. pumila* closely resemble those in other Anaeramoebae, with acetyl-CoA generated from the action of PFOR being used by a type 1C acetate:succinate CoA transferase (ASCT) to generate succinate CoA from succinate. This reaction yields the end-product acetate. Succinate-CoA, in turn, is used by succinyl-CoA synthetase (SCS) to generate ATP while regenerating succinate to continue the cycle. Additional ATP is possible to generate by the methylmalonyl-CoA pathway by a set of three enzymes, methylmalonyl-CoA epimerase, methylmalonyl-CoA mutase, and propionyl-CoA carboxylase, that convert succinate-CoA to propionate-CoA. ASCT 1B transfers the CoA moiety from propionate-CoA to succinate, yielding propionate as an end-product. The export of ATP from hydrogenosomes to the cytosol is possible using the ADP/ATP carrier family protein. Two additional mitochondrial carrier proteins were detected, but their substrates are unclear.

Interconversion of amino acids is predicted to occur in *A. pumila* MROs. We detected the presence of MRO-targeted serine hydroxymethyltransferase and the glycine cleavage chain (Figure 6) that enable the generation of CH2-THF from glycine towards serine biosynthesis. Additional enzymes involved in proline, glutamate, and threonine synthesis are also present as in other anaeramoebids^7^.

Reactive oxygen species and oxygen can damage critical components of the hydrogenosomal machinery, and protists encode several different enzymes to protect themselves. Many genes encoding these enzymes have been laterally transferred on multiple independent occasions across anaerobic eukaryote lineages ^17–19^, pointing to strong selective pressure favouring the acquisition of hydrogen-producing metabolism. We found a thioredoxin reductase (TrxR) with a predicted MTS, suggesting that H₂O₂ can be detoxified in hydrogenosomes. Oxygen detoxification can occur directly via Flavodiiron protein (Fd-FDP) or via the three enzymes: Nitrate reductase (NR), Superoxide dismutase (SOD), and Rubrerythrin (RBR) ^7,19^. OsmC can also detoxify H2O2 to water directly ^7^. Amongst these enzymes, Fd-FDP and OsmC have putative MTSs.

Collectively, the evidence indicates that the MROs in *A. pumila,* like those of *A. ignava* and *A. flamelloides,* are hydrogenosomes that produce H2, acetate, and propionate as end products.

### *Centrionella* is an anaerobic member of Legionellales and can use some end products of hydrogenosome metabolism

*C. anaeramoebae* encodes a very limited set of metabolic genes with a similar number of recognizable pathways and enzymatic reactions (74/697), as seen in *Coxiella* sp. endosymbionts found in *Amblyomma americanum* (71/623)^20^. This is in contrast to the related, but less reduced organism *Aquicella lusitana*, which encodes more than double the number of enzymes and can perform almost double the number of enzymatic reactions (159/1176). Moreover, the reduced status of the genome of the symbiont is reflected in only encoding one-third of the predicted transporters compared to those in *A. lusitana*.

#### Energy metabolism — anaerobic core

The genome lacks any genes for a conventional aerobic respiratory chain. There is no evidence of NADH dehydrogenase (Complex I), cytochrome bc1 (Complex III), or terminal oxidases. The symbiont instead appears to rely on substrate-level phosphorylation and fermentation-type pathways, which makes biological sense for an intracellular symbiont inside a host with hydrogenosomes.

*C. anaeramoebae* possesses multiple anaerobic energy-generating strategies centered on pyruvate and acetyl-CoA metabolism (Figure 7). It encodes both pyruvate formate-lyase (PFL) and its activating enzyme from a single locus, and pyruvate:ferredoxin/flavodoxin oxidoreductase (PFOR), enabling flexible conversion of pyruvate to acetyl-CoA either by generating formate (PFL) or by producing CO₂ and reduced ferredoxin (PFOR). The organism also carries acetate–CoA ligase, allowing reversible interconversion between acetate and acetyl-CoA, potentially with ATP generation, although no acetate kinase/phosphotransacetylase pair is present.

**Figure 7:**
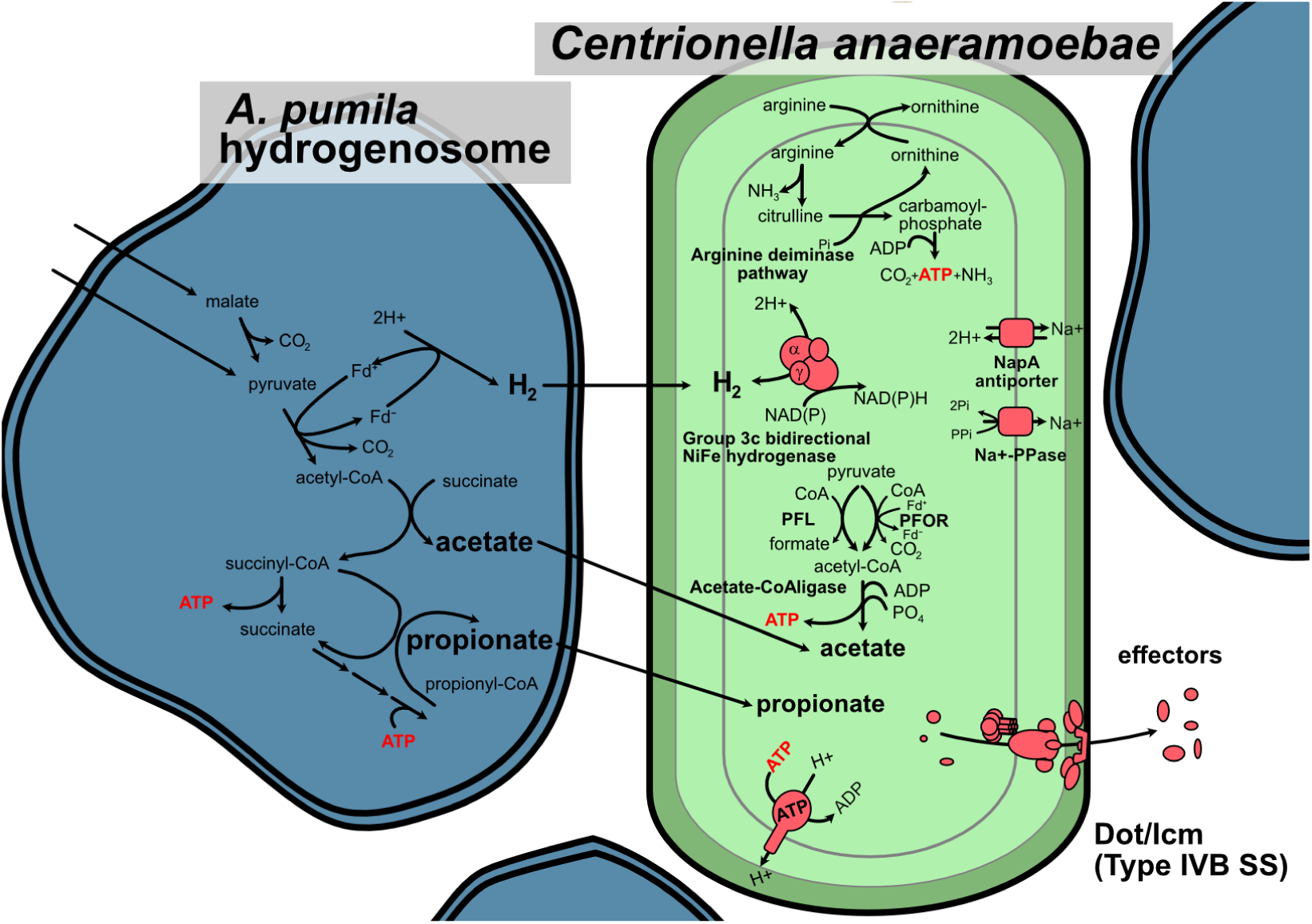
Metabolic pathways and symbiotic interactions in *Centrionella anaeramoebae.* Proposed metabolic interactions between *A. pumila* hydrogenosomes (blue compartments) and *C. anaeramoebae* (light green compartment) based on genomic and transcriptomic evidence. The hydrogenosomal compartment produces H2, acetate, and propionate as end-products. Hydrogen consumption by symbionts likely relieves product inhibition of host fermentation, enhancing overall ATP yield through syntrophic metabolism. *C. anaeramoebae* encodes a type 3c bidirectional hydrogenase that may oxidize H₂ to generate NADPH reducing equivalents. PFL cleaves pyruvate to formate + acetyl-CoA anaerobically; PFOR oxidizes pyruvate to acetyl-CoA + CO₂ while reducing ferredoxin or flavodoxin. ATP synthase — complete F₀F₁ complex can operate in reverse (ATPase mode) to pump ions to generate a proton motive force. The NapA antiporter exports Na⁺ in exchange for H⁺ import, contributing to the proton motive force. The Na⁺-PPase exports Na⁺ coupled to PPi hydrolysis, generating a sodium motive force. *Centrionella* encodes the arginine deiminase pathway, an ATP-generating fermentation pathway. Protein Abbreviations: Atp, ATP synthase. PFL, pyruvate formate-lyase; PFOR, pyruvate:ferredoxin oxidoreductase. Metabolite Abbreviations: CoA, coenzyme A; NAD, nicotinamide adenine dinucleotide; NADPH, nicotinamide adenine dinucleotide phosphate; Fdred/ox, ferredoxin or flavodoxin reduced/oxidized; ATP, adenosine triphosphate; Bold text indicates end-products and key metabolites. The Dot/Icm Type IVB and other secretion systems (only one shown) may facilitate metabolite exchange and symbiont maintenance.

The symbiont relies heavily on arginine-derived ATP generation, supported by a complete arginine deiminase (ADI) pathway encoded in a single operon together with a putative ArcD arginine/ornithine antiporter. Through the sequential activities of arginine deiminase, ornithine carbamoyltransferase, and carbamate kinase, the ADI pathway converts arginine into ornithine, ammonia, and carbamoyl phosphate, yielding one ATP per arginine, a crucial energy source in its anaerobic, host-associated niche. A second operon containing ArgR, argininosuccinate synthase, and argininosuccinate lyase suggests additional arginine regeneration from citrulline and L-aspartate, further reinforcing ATP production. The liberated ammonia may contribute to intracellular pH homeostasis, while aspartate–ammonia ligase (AsnA) may aid nitrogen balance by incorporating excess ammonia into L-asparagine. These ADI-associated genes are broadly distributed in Gammaproteobacteria, but some of the closest homologs are found in anaerobic obligate intracellular lineages such as *Parachlamydiales* and *Anoxychlamydiales*.

Ion homeostasis is maintained by two independent Na⁺ extrusion systems: a NapA-type Na⁺/H⁺ antiporter (CPA2 family) and a sodium-translocating pyrophosphatase (Na⁺-PPase). The Na⁺-PPase captures the free energy of PPi hydrolysis — a byproduct of substrate-level phosphorylation and biosynthetic reactions — to pump Na⁺ out of the cytoplasm, generating a sodium motive force. The NapA antiporter exports Na⁺ in exchange for H⁺ import, driven by the membrane potential, maintaining low cytoplasmic Na⁺ (Figure 7). The genome encodes a complete F₀F₁ ATP synthase (AtpA–I) whose c-subunit carries a standard DCCD-binding carboxylate with no Na⁺-coordinating residues, confirming proton coupling. In the absence of a respiratory electron transport chain, the ATP synthase likely operates in the hydrolytic direction, consuming ATP generated by substrate- level phosphorylation to pump H⁺ out of the cytoplasm and establish the membrane potential (ΔΨ) needed to drive NapA-mediated Na⁺ export and secondary solute transport ^21^. This arrangement — in which the F₁F₀ ATPase functions as a primary H⁺ pump rather than an ATP generator — is well established in fermentative bacteria that lack respiratory chains (Figure 7) ^22^. The phylogenetic origin of the antiporter is mixed, with many of the closest database hits being from Deltaproteobacteria, suggesting it derives from LGT. Curiously, the Na⁺/H⁺ antiporter seen in Anoxychlamydiales was also derived via LGT from Deltaproteobacteria ^5^. Together, these systems highlight a metabolic strategy centred on ion homeostasis and membrane energisation under oxygen-limited conditions, powered entirely by substrate-level phosphorylation.

For redox balance, *C. anaeramoebae* encodes a [NiFe]-hydrogenase whose catalytic subunits (FrhA, FrhG) are of archaeal Group 3c (F₄₂₀-reducing) origin, but the F₄₂₀-binding beta subunit (FrhB) is absent. Instead, the operon includes a cytochrome-c₃-type electron transfer subunit with FAD-binding and iron-sulfur domains, paired with a dedicated 4Fe-4S ferredoxin, indicating that the complex has been rewired for NAD(P)H- or ferredoxin-dependent H₂ oxidation. This reducing power feeds several NADPH-dependent enzymes, including glutamate dehydrogenase, succinate semialdehyde dehydrogenase, and enzymes of tetrahydrofolate-linked one-carbon metabolism. NAD⁺ regeneration was mediated by a soluble NADH-dependent oxidoreductase lacking membrane association and proton-pumping activity. Based on BLASTP similarity searches, the protein has the most similar homologs predominantly from Deltaproteobacteria, consistent with acquisition through lateral gene transfer. Reduced ferredoxin or flavodoxin generated by PFOR may be transferred to an HdrA-like ferredoxin and two WrbA-like flavodoxins, one of which is closely related to homologs in Desulfobacterales. Phylogenetic analyses indicate that the donor lineage might be closely related to *Desulfofustis* sp. (Desulfobulbales), sulfate-reducing strict anaerobes that often inhabit marine environments (Figure S4). The hydrogenase operon also encodes full maturation machinery — a hydrogenase maturation protease (HybD), nickel metallochaperone (HypA), nickel incorporation GTPase (HypB), and an unusual HydE–HypC fusion protein — as well as an ankyrin repeat protein inserted between the maturation and catalytic genes, potentially linking hydrogenase function to host cell interactions. Whether the [NiFe]-hydrogenase can also operate in the reductive direction — evolving H₂ from reduced ferredoxin as an electron sink when NADPH demand is met — remains to be determined. Acetyl-CoA produced by PFL or PFOR can be converted to ATP and acetate via acetate–CoA ligase (ADP-forming), providing an additional substrate-level phosphorylation route.

#### Carbon metabolism – heavily reduced glycolysis

The organism possesses only a partial lower glycolytic pathway, encoding pyruvate kinase, enolase, and a 2,3-bisphosphoglycerate-independent phosphoglycerate mutase. These enzymes enable conversion of 3-phosphoglycerate to pyruvate, providing substrate for pyruvate PFL and PFOR (Figure 7). Upstream glycolysis is absent, as key enzymes—phosphofructokinase, fructose-bisphosphate aldolase, glyceraldehyde-3-phosphate dehydrogenase, phosphoglycerate kinase, and phosphoglucose isomerase—are missing. No components of the tricarboxylic acid cycle were detected. We did not find any obvious metabolic connection to the use of propionate produced by *A. pumila* hydrogenosomes.

Given the lack of both classical glycolysis and the TCA cycle, central carbon metabolism is markedly reduced. The organism therefore likely depends on host-derived organic substrates, including amino acids, fatty acids, and acetyl-CoA precursors, rather than *de novo* carbohydrate catabolism. The mevalonate pathway is intact, with both HMG-CoA synthase and HMG-CoA reductase present. While unusual in bacteria, this pathway is retained in some Legionellales and supports isoprenoid biosynthesis.

#### Amino acid metabolism

Amino acid biosynthetic capacity in *C. anaeramoebae* is markedly reduced, consistent with its overall genomic reduction and likely host dependence. The genome encodes only a small set of enzymes associated with nitrogen assimilation and limited amino-acid interconversion. These include an NADP-dependent glutamate dehydrogenase, enabling reductive amination of α-ketoglutarate to glutamate, a central reaction in cellular nitrogen metabolism. A glutamine-independent asparagine synthetase (aspartate–ammonia ligase) catalyzes the ATP-dependent conversion of aspartate to asparagine, providing a secondary route for ammonia assimilation. In addition, serine hydroxymethyltransferase facilitates serine–glycine interconversion and contributes to one-carbon metabolism. Beyond these limited functions, most canonical amino acid biosynthetic pathways are absent, indicating that the bacterium is reliant on host-derived amino acids—a common feature of intracellular symbionts.

#### Retention of core fatty acid and phospholipid biosynthesis

*C. anaeramoebae* retains a truncated but functional type II fatty acid synthesis (FAS II) system. The elongation cycle is intact — FabG, FabZ, FabI, and FabB/FabF are all present — and the pathway initiator FabH is annotated, indicating that de novo fatty acid synthesis can proceed from malonyl-ACP. Downstream, PlsX, PlsY, and phosphatidate cytidylyltransferase support phospholipid assembly. The retention of this complete biosynthetic route from acyl chain initiation through to membrane lipid production is notable given the otherwise extreme genome reduction and indicates that *C. anaeramoebae* maintains its own membrane independently of the host — a capacity important for intracellular survival outside a host-derived vacuole.

#### Vitamins and cofactors

Cofactor biosynthesis in *C. anaeramoebae* is highly reduced, consistent with its overall genomic streamlining and similarity to *Coxiella*-like endosymbionts in arthropods ^20^. Only a limited set of short biosynthetic pathways is retained; Coenzyme A can be synthesized via the pantetheine shunt through the sequential activities of pantothenate kinase (type III), phosphopantetheine adenylyltransferase, and dephospho-CoA kinase. Elements of folate and one-carbon metabolism persist, including dihydrofolate reductase, thymidylate synthase, serine hydroxymethyltransferase, methylenetetrahydrofolate dehydrogenase/methenyltetrahydrofolate cyclohydrolase, and 5-formyltetrahydrofolate cyclo-ligase. The organism further retains two steps of the S-adenosyl-L-methionine (SAM) cycle—S-adenosylmethionine synthetase and 5′-methylthioadenosine/S-adenosylhomocysteine nucleosidase. Riboflavin can be converted to FMN and FAD via a bifunctional riboflavin kinase/FAD synthase (RibF), although no other flavin biosynthesis enzymes are present. Other cofactor pathways are highly fragmented: only a single enzyme of biotin metabolism (BioF) and the biotin–[acetyl-CoA-carboxylase] ligase (BioA2) were identified, and most pathways for folate, pterins, lipoic acid, pyridoxal-5′-phosphate, and thiamine appear absent. A limited siroheme/cobalamin branch is retained through a uroporphyrinogen-III C-methyltransferase, together with a cobalamin-binding protein. The organism is capable of converting cyanocobalamin to adenosylcobalamin via ATP:cob(I)alamin adenosyltransferase, and the presence of the BtuF cobalamin-binding protein suggests cobalamin import through a BtuCDF-type ABC transporter. Additional retained functions include sulfate activation for cysteine biosynthesis via adenylyl-sulfate kinase and sulfate adenylyltransferase (CysD), and the essential iron–sulfur cluster assembly machinery (IscU and IscS), required for the maturation of numerous Fe–S proteins, including hydrogenases. Overall, the sparse cofactor repertoire indicates strong metabolic dependence on the host, with only minimal pathways preserved for essential cofactor maintenance.

Taken together, *C. anaeramoebae* one of the first described anaerobic members of Legionellales. Its metabolism — centered on substrate-level phosphorylation, arginine fermentation, and hydrogen oxidation — closely parallels the strategies independently evolved in the Anoxychlamydiales ^5^, suggesting convergent solutions to intracellular life in oxygen-depleted environments.

### *Centrionella* maintains a large capacity to interact with its host

As expected for members of the Legionellales clade, *C. anaeramoebae* encodes a complete Dot/Icm type IVB secretion system, including IcmB, IcmD, IcmH/DotU, IcmJ/DotN, IcmQ, IcmV, IcmW, DotD, and the DotB ATPase, which together constitute the hallmark effector-delivery machinery required for intracellular niche establishment and host manipulation (Figure 3, Track 5, Figure S5). The operon is distributed across multiple genomic loci, and several components possess recognizable Sec signal peptides, consistent with periplasmic targeting before assembly. In addition to the type IVB system, the genome encodes a type II secretion system, supporting secretion of pili and extracellular proteins, as well as a type IV pilus system (PilQ, PilT, PilV) implicated in attachment and limited motility (Figure S5). A partial type VI secretion system is also present—containing TssF, TssG, TssK, TssJ, and an Hcp effector—which may mediate interbacterial interactions or host-associated functions (Figure S5). Core protein translocation capabilities are supported by a complete Sec translocon (SecABDEFGY), enabling general secretion across the inner membrane (Figure S5). Collectively, these secretion and translocation systems reflect a sophisticated host-interaction repertoire characteristic of Legionellales and underscore the organism’s adaptation to an intracellular lifestyle.

The *C. anaeramoebae* chromosome encodes eleven intact toxin–antitoxin (TA) systems — 18 type II pairs (including three higA–higB and several relB-dependent systems) and 4 type IV pairs (sdenT and abiEi types). In addition, 50 candidate anti-phage defense systems were detected, including two SoFic loci. Many of these defense-associated genes belong to the expanded gene families that are a distinctive feature of the *C. anaeramoebae* genome. Because these families may reflect selective pressures linked to the symbiont’s intracellular lifestyle, we characterized them systematically using two complementary clustering approaches:

1. The first analysis used CLANS to cluster the proteome based on all-against-all BLAST pairwise similarities (Figure 8A). Manual annotation of notable gene clusters revealed eight gene clusters associated with mobile elements of various types (recombinases, transposases, six types of IS-elements). The other large gene families consisted of proteins with Cadherin-like domains (n=33), ATP-binding proteins (n=40 and n=6), Ankyrin-repeat proteins (n=41), Protein kinases (n=24), and AAA-type ATPases (n=14 and n=6). The Ankyrin-repeat proteins, Protein kinases, one of the ATP-binding proteins, and one of the AAA-family ATPase clusters form a large, connected network, likely indicating domain sharing (Figure 8A). A cluster of putative non-ribosomal peptide synthases was also detected.
2. The second clustering method used OrthoVenn to cluster gene families in *Centrionella* against *Candidatus* Carsonella ruddii, a highly reduced Gammaproteobacterial genome lacking secretion systems and thus expected to lack effectors. There were 127 protein clusters and 786 singletons in Centrionella. The clustering revealed 78 unique gene families with a total of 156 proteins with more than 2 members (Figure S6). Multi-member clusters (n≥3) were plotted on the CLANS-generated chart showing good agreement between gene family prediction methods (Figure 8B). As seen above, amongst the largest gene families, there were several distinct families that have similarity to eukaryotic-like domains (Cadherin-like domains, Ankyrin-repeat domains, WD40-repeat domains) as well as AAA-family ATPase or ATP-binding proteins (Figure S7). The single largest gene family was again the 33 Cadherin-like domain proteins (CHDL). CHDLs have been detected in some bacteria, such as proteobacteria and cyanobacteria ^23^. BLASTP searches place the *C. anaeramoebae* CHDLs closest to bacterial homologs from Gammaproteobacteria, Cyanobacteria, and Chlamydiota, with no single donor lineage dominating the top hits. A CHDL protein has been implicated in host adherence and cytotoxicity in the protist parasite *Trichomonas vaginalis,* where it is surface-localized and it induces both host-parasite and parasite-parasite aggregation ^24^. The second-largest cluster comprises 25 Ankyrin repeat proteins (ARPs). These proteins generally show the highest similarity to eukaryotes, but of diverse phylogenetic origin, suggesting a dynamic evolutionary history. Interestingly, some ARPs show weak similarity to trichomonad and anaeramoebid ARPs.

**Figure 8:**
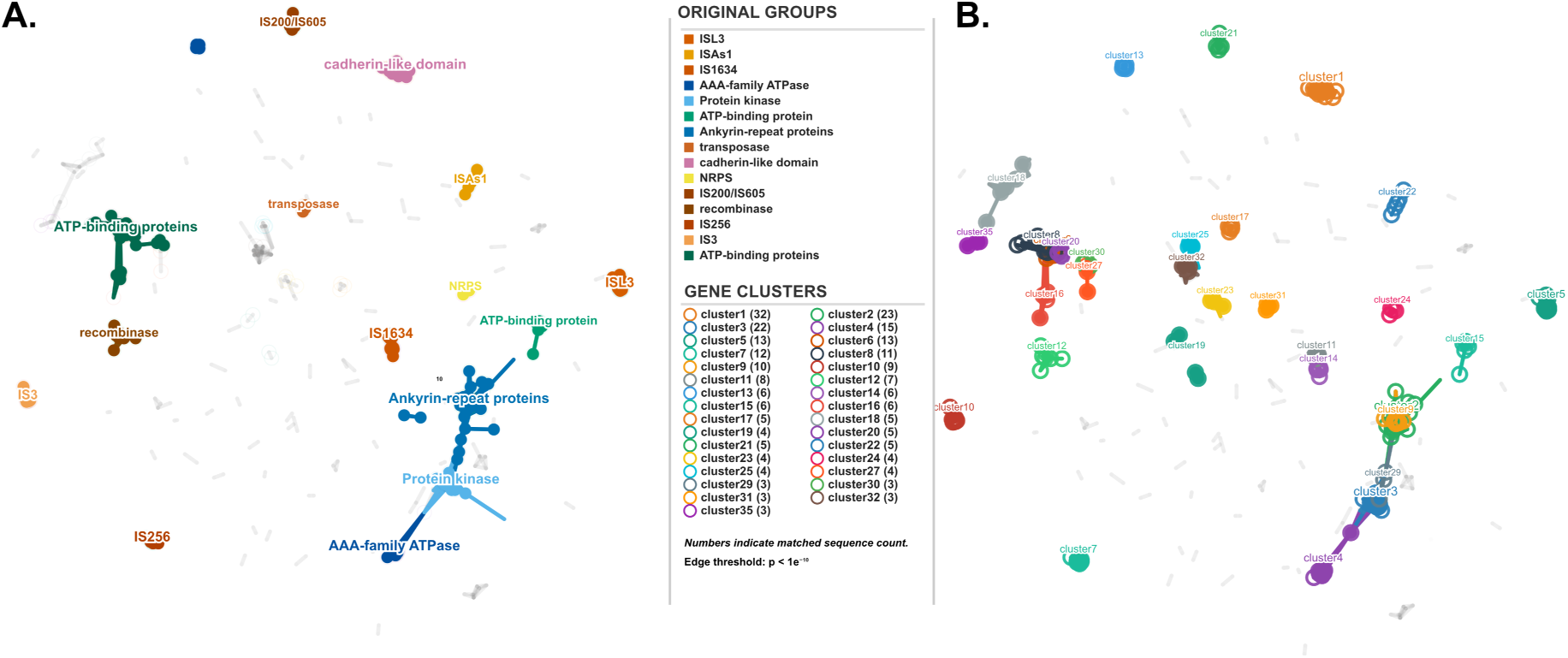
The *Centrionella anaeramoebae* proteome has large gene families with eukaryotic-like domains. The *C. anaeramoebae* proteome was clustered using CLANS v.29.05.2012^5^ at an e-value limit of 9.2E-5 to cluster protein families. **(A)** shows 15 manually annotated protein clusters, Ankyrin-repeat proteins — 41 seqs (blue), ATP-binding proteins — 40 seqs (green), cadherin-like domain — 33 seqs (pink), Protein kinase — 24 seqs (light blue), AAA-family ATPase — 20 seqs (blue), IS1634 — 14 seqs (red, transposon), IS256 — 14 seqs (red, transposon), ISL3 — 13 seqs (red, transposon), IS3 — 10 seqs (orange, transposon), recombinase — 8 seqs (red), ATP-binding protein — 6 seqs (light green), IS200/IS605 — 5 seqs (red), ISAs1 — 5 seqs (red), transposase — 5 seqs (red), NRPS — 4 seqs (yellow, non-ribosomal peptide synthetase) and **(B)** shows an overlay of unique gene family clusters with more than 3 members clustered via OrthoVenn3^6^.

Beyond the large families, several smaller clusters encode multidomain proteins with eukaryotic-like architectures. For example, the three members of cluster 29 each carry a putative nucleotidyltransferase/adenylyltransferase domain fused to varying combinations of ankyrin repeats, WD40 repeats, and signalling-associated domains (Rab, Ran, AMN1) (Figure 8B). The modular domain organization of these proteins is consistent with multifunctional effectors that could engage different host targets.

The prevalence of eukaryotic-like domains in the large gene families is a hallmark of effector proteins in other Legionellales, raising the possibility that many of these proteins are translocated into the host via the Dot/Icm system. To assess this, we used the S4TE prediction server ^25^ to identify candidate type IVB secretion effectors across the *C. anaeramoebae* proteome. The analysis identified 184 putative effectors, encompassing 414,900 bp of coding sequence — 27.3% of the genome and 31.0% of all CDS bases (Figure S6). Many correspond to members of the large gene families described above, including all three cluster 29 proteins (Figure 9A). The predicted effectors are distributed across the chromosome with local clusters apparent at several loci (Figure 3, Track 6), and their mean gene length (2,255 bp) substantially exceeds the bacterial average, consistent with the multidomain architectures observed in the family-level analysis.

**Figure 9:**
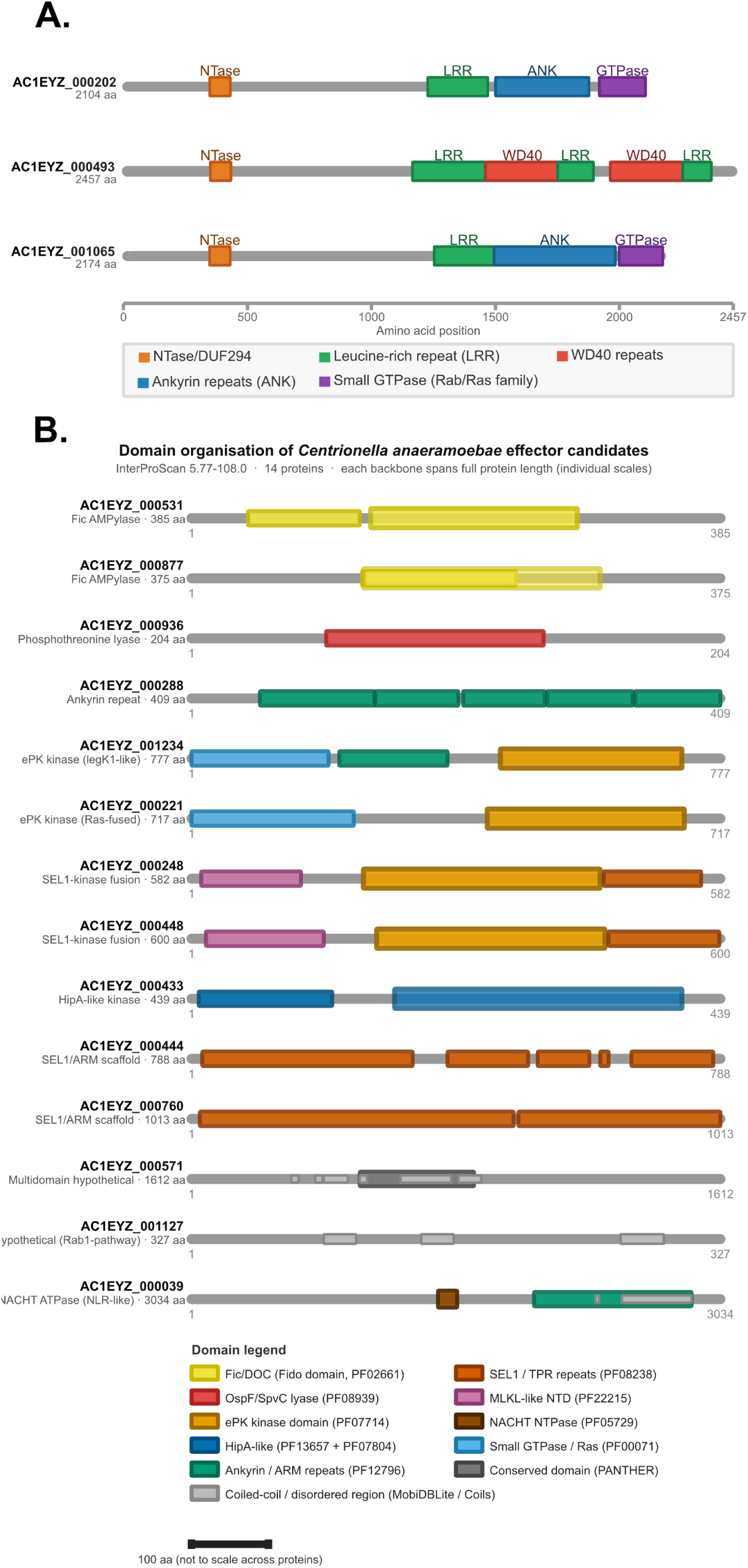
Domain organization of effector candidate proteins from *Centrionella anaeramoebae.* **(A)** Protein domain architecture of the three large multi-domain proteins with eukaryote-like domain of protein cluster 29. **(B)** Domain organization of selected *C. anaeramoebae* effector candidates. Horizontal lines represent full-length protein sequences drawn at an individual scale (each backbone = full protein length). Colored rectangles indicate domain boundaries predicted by InterProScan 5.77-108.0 against Pfam, SMART, ProSiteProfiles, Gene3D, PANTHER, and SUPERFAMILY databases. Locus tags use PGAP annotation identifiers (AC1EYZ_XXXXXX). Protein lengths (aa) are given in the protein label. ANK, ankyrin repeat; ARM, armadillo repeat; ePK, eukaryotic protein kinase; MAPK, mitogen-activated protein kinase; MLKL, mixed-lineage kinase domain-like; NACHT, domain present in NAIP, CIITA, HET-E and TP1; NLR, NOD-like receptor; SEL1, Sel1-like repeat; TPR, tetratricopeptide repeat.

Among the predicted effectors, several multidomain proteins stand out for their unusual domain architectures (Figure 9B, Figure S7). AC1EYZ_000221, for example, combines a Rho-family small GTPase domain with ankyrin repeats and a C-terminal eukaryotic-type Ser/Thr kinase — a domain combination not described in other Legionellales. Two neighboring genes encode related proteins with shared but distinct architectures: AC1EYZ_001234 carries an N-terminal small GTPase domain, ankyrin repeats, and a eukaryotic-like protein kinase, while AC1EYZ_001232 retains the GTPase and ankyrin repeat domains but lacks the kinase, with only a short C-terminal fragment (26 aa) preserved (Figure 10B). The small GTPase domains in these proteins show high similarity to Rac1 homologs in anaeramoebids, and phylogenetic analysis recovers them as most closely related to a Rac1 protein in *A. pumila* (tig00005939_segment0.cds333.1), which is itself present as four identical tandem copies (Figure 10A). Pairwise alignments between the *Centrionella* and *A. pumila* proteins reveal 70–75% identity across homologous regions, supporting horizontal acquisition of a host Rac1 gene by *Centrionella*, followed by domain shuffling and gene duplication to generate the observed variants. An additional effector candidate (AC1EYZ_000939) carries a BEACH domain with the highest similarity to an *A. flamelloides* homolog, suggesting that horizontal acquisition of host-derived domains may extend beyond the Rac1 family. Both Rac1-domain proteins are expressed at the RNA level, but whether they are translocated into the host cytoplasm and what functions they perform remain open questions. Rac1 is a central regulator of eukaryotic cytoskeletal dynamics and a frequent target of pathogen interference, making the direct acquisition of host Rac1 genes — rather than the production of mimicking effectors — a notable departure from described strategies of host manipulation.

**Figure 10:**
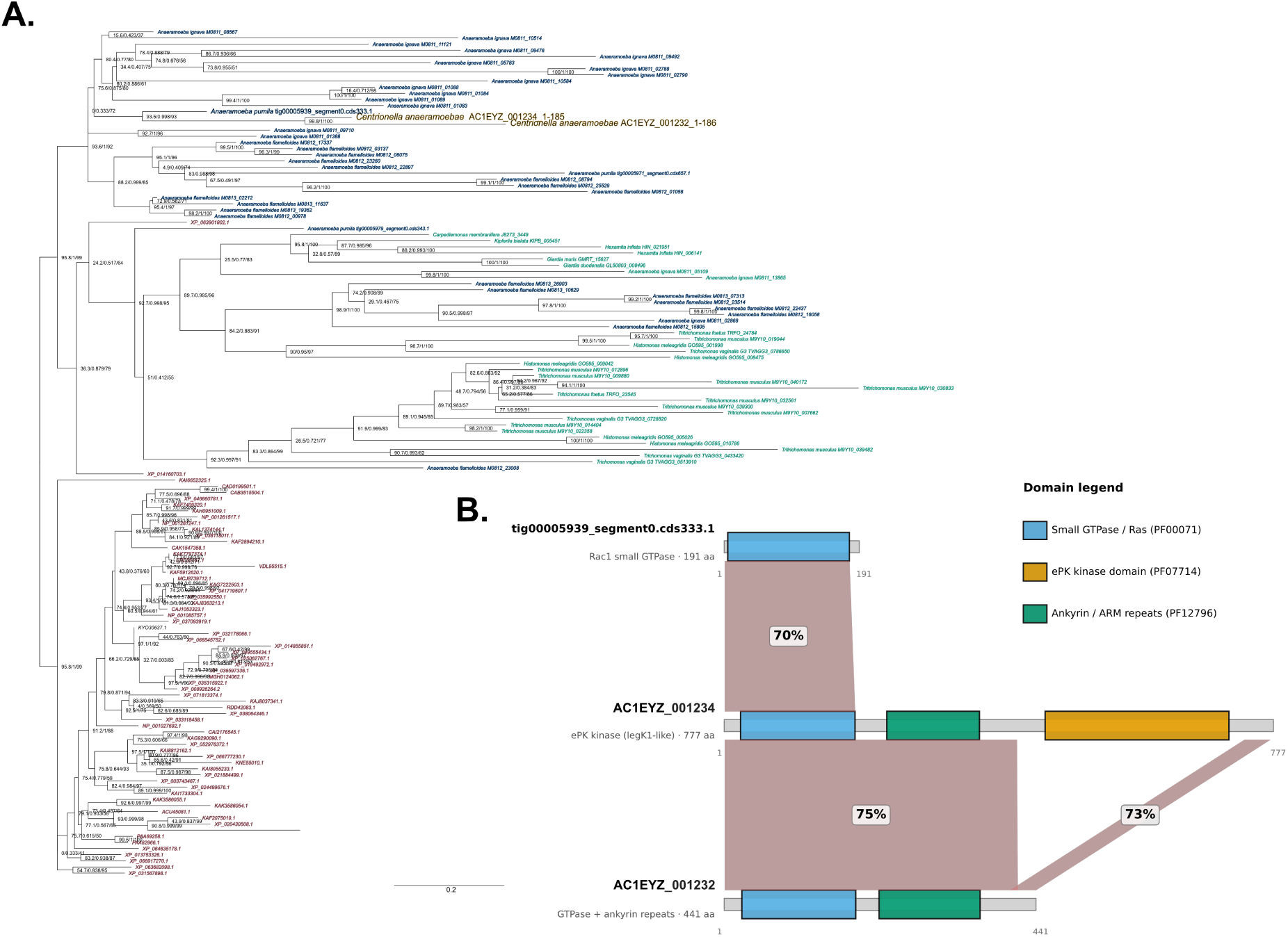
Domain architecture and sequence similarity of *Centrionella anaeramoebae* proteins acquired through horizontal gene transfer from Anaeramoebae. (A) Phylogenetic analysis of 138 Rac1 domain proteins (191 sites, 182 informative) in Anaeramoebae (blue), *C. anaeramoebae* (AC1EYZ_001234 (1-185) and AC1EYZ_001232 (1-186)) (gold), Metamonada (green), and other eukaryotes (red). The phylogeny was estimated using IQ-TREE 3.1.0, with ModelFinder selecting LG+R5 as the optimal substitution model. The support values within parentheses are SH-aLRT support (%) / aBayes support / 1000 ultrafast bootstrap support (%). **(B)** The top protein from *A. pumila* (tig00005939_segment0.cds333.1, 191 amino acids), the middle protein (AC1EYZ_001234, 777 amino acids) is a fusion protein containing N-terminal GTPase, central ankyrin repeat, and C-terminal serine/threonine kinase domains. The bottom protein (AC1EYZ_001232, 441 amino acids) contains GTPase and ankyrin repeat domains but lacks the kinase domain, suggesting either gene truncation or domain loss. The proteins are shown as horizontal tracks with their predicted domain architectures based on InterProScan analysis. Colored rectangles represent functional domains: blue indicates small GTPase/Ras domains (PF00071), orange shows ePK kinase domains (PF07714), and green depicts ankyrin repeat domains (PF12796). Gray ribbons connecting the proteins indicate regions of sequence similarity identified by BLASTP analysis, with percentages showing amino acid identity. Scale bars indicate protein length in amino acids.

*Centrionella* encodes two Fic domain proteins capable of covalent AMPylation of target proteins. Fic-mediated AMPylation is a well-characterized virulence strategy in intracellular pathogens — IbpA from *Histophilus somni*, for example, AMPylates Rho GTPases to collapse the host actin cytoskeleton ^26^. Both *Centrionella* Fic proteins were additionally flagged as candidate SoFic anti-phage systems, and future work is needed to distinguish their roles in host manipulation from those in phage defense.

### *Desulfobacter* sp. LANTAAN and tripartite syntrophy

The 16S rDNA analysis revealed a *Desulfobacter* sp. in the *A. pumila* culture that was enriched in amoebae-containing fractions harvested by cold-shock but partitioned to the pellet fraction upon Histopaque separation (Figure 2A). FISH with the Delta495a probe identified long rod-shaped bacteria in close proximity to host cells but not located within them (Figure 11A). The fully assembled genome of the *Desulfobacter* sp. is a single circular contig of 7.47 Mbp encoding 6,361 CDSs (Figure S8) — among the largest reported for *Desulfobacter* sensu stricto, consistent with a free-living, metabolically versatile lifestyle. A detailed metabolic reconstruction is provided in Supplementary Note 1. Phylogenetic analysis places *Desulfobacter* sp. LANTAAN close to *Desulfobacter* sp. isolates associated with *A. ignava* ^8^ (Figure 11B), indicating a recurrent association between Anaeramoebidae and this clade of sulfate reducers.

**Figure 11:**
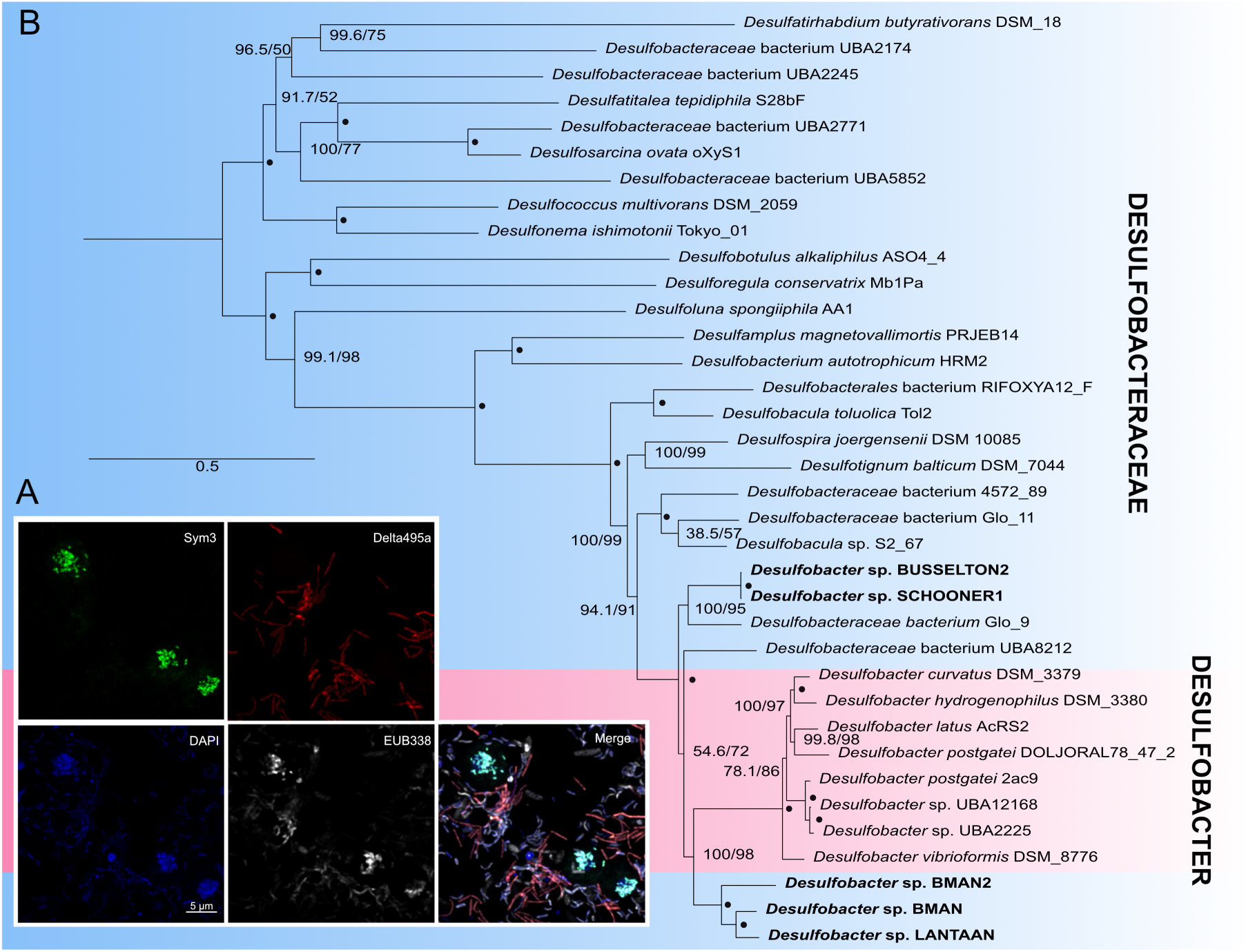
FISH of *A. pumila* consortia and phylogenomic analysis of *Desulfobacter* sp. LANTAAN. **(A)** FISH of *A. pumila* consortia blue, green, white, and red signals represent DAPI, *Centrionella*-specific (Sym3) probe, universal bacterial (EUB338) probe, and Deltaproteobacteria (Delta495a) probe, respectively. **(B)** The maximum-likelihood (ML) tree was constructed from concatenated data for 94 orthologous proteins using the LG+C60+F+G model in IQ-TREE. *Desulfobacter* sp. in *A. pumila* consortia and symbionts in *A. flamelloides* BUSSELTON2 and SCHOONER1, *A. ignava* BMAN/BMAN2 are marked with bold font. Solid circles at branch nodes indicate full confidence values, while numerical values represent standard confidence values. The scale bar indicates a substitution rate of 20%.

The metabolic repertoire of *Desulfobacter* sp. LANTAAN fits well with a putative syntrophic partnership with *A. pumila*. The organism can oxidize all major hydrogenosomal end-products — H₂, acetate, and propionate — via dissimilatory sulfate reduction, with the Wood–Ljungdahl pathway serving both as the oxidative route for acetate catabolism and as a reductive CO₂-fixation pathway. This complementarity relieves product inhibition of host fermentation while providing electron donors for sulfate reduction. The genome also encodes a complete cobalamin biosynthesis pathway (53 loci) and dedicated B₁₂ transporters. *A. pumila* possesses a corresponding utilization pathway — LMBRD1, MMAB, MCEE, MUT, and MetH — indicating adenosylcobalamin dependence in propionyl-CoA catabolism and methionine synthesis, consistent with B₁₂ provisioning by *Desulfobacter*. The B₁₂-dependent enzymes unique to *A. ignava* and *A. flamelloides* ^8^ were not found in *A. pumila*, suggesting lineage-specific variation in the scope of this nutritional interdependence.

Together with *C. anaeramoebae*, *Desulfobacter* sp. LANTAAN appears to form a tripartite syntrophic consortium centred on the end-products of host hydrogenosome metabolism. *C. anaeramoebae* can oxidize H₂ via its bidirectional [NiFe]-hydrogenase and activate host-derived acetate through acetate–CoA ligase, while *Desulfobacter* sp. LANTAAN performs the bulk of sulfate-dependent hydrogen and acetate consumption. Propionate utilization by *C. anaeramoebae* seems unlikely, as no canonical methylmalonyl-CoA pathway is encoded. The partitioning of syntrophic roles between the two bacterial partners — with *Centrionella* as a metabolically reduced intracellular consumer and *Desulfobacter* as a versatile extracellular scavenger — parallels the community-level metabolic organization recently described in anaerobic breviate protist consortia, where protist fitness depends on the collective metabolic function of associated bacteria rather than on any single partner ^27^.

## Discussion

*Centrionella anaeramoebae* represents an anaerobic member of Legionellales, a lineage otherwise comprising aerobic intracellular pathogens, typified by *Legionella pneumophila* ^28^. Ca. Centrionella is, to our knowledge, the first genomically characterised anaerobic member of Legionellales from a marine environments. Its genome encodes an entirely anaerobic metabolism built on substrate-level phosphorylation, arginine fermentation, and hydrogen oxidation — a metabolic architecture that has replaced aerobic respiration entirely. This transition parallels the independent adaptation of Anoxychlamydiales from aerobic Chlamydiae ancestors to anaerobic symbionts ^5^. The convergence extends to specific metabolic solutions: both lineages have arrived at arginine deiminase pathways for ATP generation, and both have acquired Na⁺/H⁺ antiporter systems via horizontal gene transfer from Deltaproteobacteria ^5^. The retention of hydrogen-metabolizing enzymes in *C. anaeramoebae* despite extensive genome reduction points to the ecological importance of syntrophic hydrogen transfer in these partnerships. That phylogenetically distant obligate intracellular lineages converge on similar metabolic strategies suggests a limited set of viable solutions for anaerobic intracellular life, constrained by the end-products available from host hydrogenosome metabolism.

The retention of a complete Dot/Icm type IVB secretion system alongside expanded gene families carrying eukaryotic-like domains indicates that host manipulation remains central to the *Centrionella* lifestyle, despite the apparent shift from pathogenesis toward mutualism. The Dot/Icm system is essential for *Legionella* pathogenesis, mediating establishment of replicative vacuoles and host cell manipulation ^29^. In *C. anaeramoebae*, this machinery is presumably repurposed for symbiont maintenance and host interaction. The most striking finding in this context is the presence of eukaryotic Rac1-like GTPase genes acquired from the host. Eukaryote-to-prokaryote gene transfer is considered rare and typically involves metabolic genes rather than core cellular machinery ^30,31^. Intracellular pathogens commonly manipulate host Rho GTPases through effectors that mimic GEF activity, such as *Salmonella* SopE ^32^ or *Shigella* IpgB1 ^33^, or through indirect modulation via upstream regulators like *Legionella* LepB ^34^. The *Centrionella* case differs: rather than encoding effectors that act on host GTPases, the symbiont has acquired the GTPase genes themselves, followed by domain shuffling and duplication into distinct domain architectures. Whether these proteins function as effectors trafficked into the host cytoplasm, or serve endogenous functions within the bacterium, remains to be determined experimentally.

The metabolic integration between *A. pumila* hydrogenosomes, *Centrionella*, and *Desulfobacter* sp. LANTAAN could constitute a tripartite syntrophic consortium (Figure 12). Recent work on anaerobic breviate protists has shown that protist fitness depends on the metabolic capabilities of associated hydrogen-consuming bacteria rather than their taxonomic identity ^27^. The *A. pumila* system may exhibit a similar organization, with *Centrionella* consuming H₂ via its bidirectional [NiFe]-hydrogenase, while *Desulfobacter* sp. LANTAAN performs dissimilatory sulfate reduction using hydrogenosomal end-products. That *Desulfobacter* sp. LANTAAN is phylogenetically related to symbionts of other *Anaeramoeba* species ^8^, and encodes complete vitamin B₁₂ biosynthesis pathways corresponding to B₁₂-dependent enzymes in *A. pumila*, indicates nutritional interdependencies beyond hydrogen scavenging. Similar B₁₂-based interdependencies and sulfide provisioning have been documented in breviate consortia ^27^.

**Figure 12:**
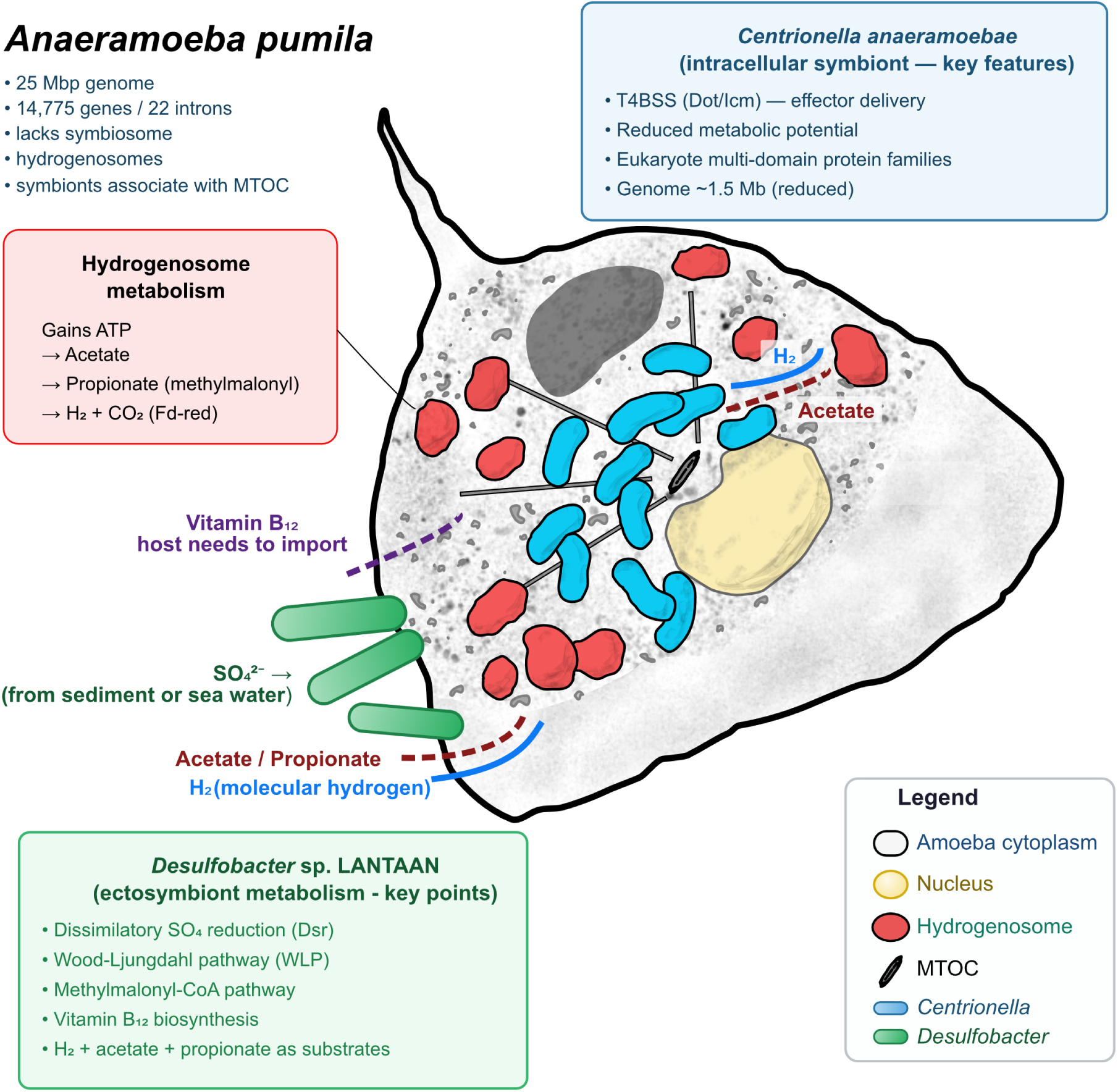
*Anaeramoeba pumila* consortia syntrophic interactions and features. Putative syntrophic interactions present in the *A. pumila* consortia. *Centrionella anaeramoebae* infect *A. pumila* and are closely associated with the microtubule-organizing centre (MTOC). It encodes several secretion systems, including T4BSS (Dot/Icm) and large gene families of proteins with eukaryote-like domains similar to effectors described in other members of the Legionellales. The hydrogenosomes of *A. pumila* produce molecular hydrogen (H2), acetate, and propionate. H2 and acetate are likely used by *C. anaeramoebae*. All three end-products of hydrogenosomal metabolism are predicted to be used by the sulfate-reducer *Desulfobacter* sp. LANTAAN. *Desulfobacter* sp. LANTAAN encodes a complete cobalamin biosynthesis pathway and might provide Vitamin B12 to the consortia.

The absence of symbiosomes in *A. pumila*, despite its phylogenetic position emerging from within a clade of symbiosome-bearing taxa, indicates secondary loss. Two non-exclusive explanations merit consideration. First, the small cell size of *A. pumila* may permit diffusive clearance of H₂ without requiring the close apposition of hydrogenosomes and symbionts that the symbiosome provides in larger *Anaeramoeba* species. Second, *Centrionella* effectors — including the HGT-derived Rac1/Rho-domain proteins — could actively suppress symbiosome biogenesis to favor the cytosolic niche. The localization of *Centrionella* to the MTOC rather than a membrane compartment is consistent with cytoskeletal anchoring strategies employed by other endosymbionts, notably *Wolbachia* in nematodes ^35^, and the large CHDL protein family in *Centrionella* may contribute to MTOC positioning and co-segregation during cell division. This organizational strategy — cytoskeletal association without a dedicated membrane compartment — may represent the primary mechanism for intracellular positioning in this system. Introducing *Centrionella* into symbiosome-bearing *Anaeramoeba* isolates would provide a direct test of whether the effector repertoire is sufficient to suppress symbiosome formation.

### Description of *Candidatus* Centrionella anaeramoebae gen. nov., sp. nov

***Centrionella anaeramoebae*** (Cen.tri.o.ne’lla. **L. neut. n.** centrum, center; **Gr. dim. suff.** -ion, denoting smallness; **N.L. fem. dim. suff.** -nella, denoting smallness; **N.L. fem. n.** *Centrionella*, small [organism] at the center, pertaining to the close affiliation of the organism to the microtubule-organizing centre (MTOC) of its native host cell; an.ae.ra.moe’bae, **N.L. gen. n.** anaeramoebae, of *Anaeramoeba*, referring to the host genus, pertaining to the family name Anaeramoebidae of the host species *Anaeramoeba pumila* LANTAAN.

**Classification:** Pseudomonadota; Gammaproteobacteria; “*Ca.* Centrionellales”; “*Ca.* Centrionellaceae”; “*Ca.* Centrionella”, “*Ca.* Centrionella anaeramoebae”

**Morphology:** Rod-shaped Gram-negative bacteria; 300–650 nm long and approximately 100–300 nm wide. Non-motile; no flagella or basal bodies observed by transmission electron microscopy. Cells lack a distinct pericellular membrane of host origin and are not enclosed in a vacuole or symbiosomal compartment.

**Ecology and lifestyle:** Obligate intracellular symbiont of *Anaeramoeba pumila* LANTAAN, residing in the host cytoplasm in close association with the permanent microtubule-organizing centre (MTOC) and microtubule network radiating from it. The bacteria are stably maintained through host cell cycles and are segregated during cell division. Strictly anaerobic microenvironment; no growth outside the host cell has been reported.

**Phylogenetics:** GTDB-Tk places the organism within the class Gammaproteobacteria, order “*Ca.* Centrionellales” (formerly UBA12402), family “*Ca. Centrionellaceae*” (formerly UBA12402), most closely related to the genus *Aquicella* (*Coxiellaceae*) but sufficiently divergent to constitute a new family and genus. Since this represents the first described type species of UBA12402, the order was named “*Ca.* Centrionellales”. T4BSS (Dot/Icm) components consistent with the Legionellales common ancestor T4BSS are predicted.

**Host organism:** The amoeba host *Anaeramoeba pumila* LANTAAN was isolated from marine anoxic sediments at Koh Lanta island, Thailand (7°28′40.3″ N, 99°06′20.5″ E). The host 18S rRNA gene sequence is deposited at GenBank/ENA/DDBJ under accession number OQ924551.

**Nucleotide sequence accessions:** Basis of assignment: 16S rRNA gene (GenBank/ENA/DDBJ accession number LT716083). Oligonucleotide probe Sym3 (5′-CCTCTACTGAACTCTAGTCATCTAGT-3), hybridizing at *E. coli* 16S rRNA gene position 641.

## Materials and methods

### Cultivation and enrichment of *A. pumila*

The *Anaeramoeba pumila* cells were maintained in sealed cell culture tubes (Thermo Fisher Scientific, USA) containing 10 mL of 50% SW802 medium, with weekly subculturing. The SW802 medium was prepared by mixing 1× ATCC medium (Cereal Grass Media; Ward’s Science, USA, Cat. No. 470300-680) with 1× Artificial Seawater (ASW) at a 1:1 ratio. The ASW formulation per liter is: 24.72 g NaCl, 0.67 g KCl, 1.364 g CaCl₂·2H₂O, 4.66 g MgCl₂·6H₂O, 6.29 g MgSO₄·7H₂O, and 0.18 g NaHCO₃.

For large-scale cultivation, 550 mL cell culture flasks (Greiner Bio-One, Germany, 660160) were used with 25% SW802 medium. Due to the adherent growth of cells, a combination of cold shock, physical impact, and cell scraping was employed for enrichment: First, the supernatant in the culture flask was slowly discarded, and the cells were rinsed once with 100 mL ASW. Then, 50 mL of pre-chilled (4°C) ASW was added, and the flask was placed on ice for cold shock about 10 minutes. After microscopic examination, the flask was dropped from a height of about 0.5 meters several times to detach the amoebae from the flask surface effectively. A cell scraper (Falcon™ Cell Scrapers, Thermo Fisher Scientific, USA, Cat. No. 353087) was then used to scrape the bottom of the flask to maximize cell collection. Finally, the suspension was centrifuged (500 × g, 8 minutes, 4°C) to pellet cells.

### DNA, RNA extraction, and high-throughput sequencing

Total DNA was extracted using the CTAB method ^36^and subsequently purified with the Genomic-tip 20/G kit (Qiagen, Hilden, Germany; Cat. No. 10223) to remove impurities and RNA. The purified genomic DNA (concentration: 109 ng/µL; A260/A280: 1.887; A260/A230: 2.499) was submitted to the Centre d’expertise et de services Génome Québec (Canada) for construction of a Shotgun PCR-free library and Illumina sequencing. Genomic DNA for nanopore sequencing was treated with the Short Read Eliminator Kit (Circulomics, USA, ss-100-101-01) to remove short DNA fragments, and a sequencing library was constructed with the Ligation Sequencing Kit (Oxford Nanopore Technologies, UK, Cat. No. SQK-LSK109) and Flow Cell Priming Kit (Oxford Nanopore Technologies, UK, Cat. No. EXP-FLP002). Sequencing was performed on a MinION Mk2 device controlled by MinKNOW software.

Total RNA was extracted using the TRIzol method (Ambion, USA, Lot No. 210806) ^37^. Residual DNA was removed using the TURBO DNA-free™ Kit (Thermo Fisher Scientific, USA, Cat. No. AM1907). The RNA sample was also submitted to the Centre d’expertise et de services Génome Québec (Canada) for Illumina sequencing on NovaSeq 6000 platform using a strand-specific protocol.

### Metagenome assembly and host genome extraction, gene prediction and annotation

Basecalling was first performed on the raw long-read sequencing data (fast5 files) using Guppy v.3.4.4 (Oxford Nanopore), converting them to fastq files. Adapter trimming was then conducted using Porechop v.0.2.4 (https://github.com/rrwick/Porechop). The genomic data were assembled separately using Flye v.2.7 ^38^ and Canu v.2.0 ^39^. The initial assembly results were polished using Racon v.1.4.20 (4 rounds) ^40^ and Medaka v.0.11.5 (https://github.com/nanoporetech/medaka). The short-read genomic sequencing data were quality-trimmed using Trimmomatic v.0.36 ^41^ and used with the unicycler_polish function in Unicycler v.0.4.8 ^42^ to further correct errors in the nanopore-based assemblies (5 rounds for the Flye assembly, 11 rounds for the Canu assembly). This resulted in Flye assemblies and Canu assemblies, which were compared and analyzed using QUAST v.5.0.2 ^43^.

RNA-seq data were trimmed using Trimmomatic v.0.36 ^41^ and *de novo* assembled using Trinity v.2.11.0 ^44^. Since the Canu assembly yielded a superior N50 (1,865,526 bp) compared to Flye (677,806 bp), it was selected for host genome extraction as follows: RNA-seq, Illumina, and Nanopore sequencing data were mapped to the Canu metagenomic assembly contigs using HISAT2 v.2.1.0, Bowtie2 v.2.3.0, and ngmlr v.0.2.7, respectively ^45,46^. Based on the alignment profiles from the three datasets, combined with the GC content of the contigs and BLASTn search results against the NCBI non-redundant nucleotide (nr) database, the *A. pumila* genome contigs were identified and extracted from the metagenome.

Gene prediction for the extracted host genome was performed using GlimmerHMM. Examples of introns were obtained from manual observation and added to the annotation, identifying 22 introns in 19 genes. Due to gene duplications and examples of identical sequences amongst introns, 17 unique intron sequences were used in the training set. Predicted gene structures were verified and refined using RNA-seq alignment evidence, resulting in the final host proteome. The proteome was annotated using Funannotate v.18.17 with database 2023-09-28-003158 as implemented in Galaxy Europe (2026-04-02) ^47^. The annotations were based on input from eggnog-mapper v.2.1.13 ^48^ with the eggnog 5.0.2 database ^49^ using --very-sensitive for diamond searches, InterProScan Galaxy Version 5.59-91.0+galaxy3 ^50^, and the Phobius webserver (https://phobius.sbc.su.se/, accessed 2026-04-01) ^51^. To identify mitochondrion-related organelle (MRO) proteins of *A. pumila*, the MRO proteomes of *A. flamelloides* strains BUSSELTON2 and BMAN were used as references for BLASTP searches.

Concurrently, homologous searches for nucleus-encoded mitochondrial proteins and predictions of mitochondrial targeting sequences (MTS) were conducted using TargetP, Mitoprot, and Mitofates against three databases: the NCBI nr database, the mitochondrial database (MitoDB), and the metamonad protein database (CLODB) (from Prof. Andrew J. Roger’s laboratory, Dalhousie University). Integrating the results from these analyses, the MRO proteome of *Anaeramoeba pumila* was ultimately determined.

### Non-coding RNA prediction and annotation

A total of 206 high-confidence non-coding RNA loci were annotated across the *A. pumila* genome: 110 primary rRNA loci (Infernal-trusted), 96 tRNA loci (≥2 prediction tools + 2 selenocysteine (SeC)). The SeC tRNA loci were detected exclusively by ARAGORN and included unconditionally, given their biological significance as a functionally essential class indicating active selenoprotein synthesis.

tRNAs were predicted and annotated using tRNAscan-SE v2.0.12 (flags: -E –max --gff) (^52^ and ARAGORN v1.2.41 (flags: -t -i -l -o) ^53^. The tabular output from ARAGORN was converted to gff-format using cnv_aragorn2gff.pl (https://github.com/jestill/dawgpaws/tree/master/scripts/cnv_aragorn2gff.pl). Annotations were joined, and tRNAs (≥2 tools agree) were kept, except for two SeC tRNA candidates that were solely predicted by ARAGORN. In total 94 tRNAs were annotated (20 isotypes + SeC). Ribosomal RNAs (LSU 28S rDNA, SSU 18S rDNA, 5.8S rDNA, 5S RNA) were predicted by Infernal v1.1.5 (flags: --tblout) ^54^ using the Rfam covariance models ^55^ (5S_rRNA - RF00001, 5_8S_rRNA - RF00002, SSU_rRNA_eukarya - RF01960, LSU_rRNA_eukarya - RF02543) downloaded 2026-03-28 and barrnap v0.9 (https://github.com/tseemann/barrnap) (flags: kingdom --euk). The Infernal output was processed to gff-format using the script infernal_tblout2gff.pl (https://raw.githubusercontent.com/nawrockie/jiffy-infernal-hmmer-scripts/master/infernal-tblout2gff.pl). We noticed that barrnap did not predict complete 28S rRNA loci but consistently split them into three parts. These were successfully recovered as single loci by Infernal. In total, 49 LSU (28S rRNA) genes (22 complete and 27 partial), 24 SSU (18S rRNA) genes (22 complete and 2 partial), 24 5.8S rRNA genes, and 13 5S rRNA genes were annotated. All 22 complete 45S rDNA repeats include a paired 5.8S rDNA copy. Partial 28S rDNA repeats without another rDNA terminate 19 contigs.

### Bacterial genome sequencing and analyses

#### Fractionation of *A. pumila* and bacteria

Both cold shock and Histopaque separation methods were used for the fractionation of *A. pumila* cells and bacteria. For cold shock, the *Anaeramoeba pumila* cells were enriched as described above in ‘Cultivation and enrichment of *Anaeramoeba pumila’*. The culture supernatant was centrifuged (2791 × g, 20 minutes, 4°C) to pellet planktonic bacteria, resulting in bacterial-enriched samples.

For the second method, Histopaque®-1077 (Sigma-Aldrich, USA, Lot No. RNBG5100, density 1.077 g/mL) was added to a glass centrifuge tube, sealed with parafilm, and pre-warmed to room temperature. Enriched *A. pumila* samples were slowly layered onto the Histopaque and centrifuged at room temperature (2000 × g, 20 minutes), resulting in a stratified solution: bacteria pelleted at the bottom, and amoeba cells concentrated in the intermediate layer. The intermediate amoeba-enriched layer was aspirated and examined under a microscope to ensure minimal bacterial contamination. After that, the cells were diluted with room-temperature ASW, pelleted by centrifugation (2791 × g, 2 minutes, room temperature), and the supernatant was discarded to obtain a final amoeba-enriched sample free from Histopaque. Separately, the bacterial pellet from the Histopaque centrifugation was collected after removing the supernatant, resuspended in room-temperature ASW, and examined microscopically, confirming the presence predominantly of prokaryotic bacteria. This suspension was also centrifuged (2791 × g, 2 minutes, room temperature), and the supernatant was discarded to obtain the bacterial-enriched sample.

#### DNA extraction and 16S rDNA sequencing

For all the samples obtained from the two enrichment methods above, each was resuspended in CD1 solution (from the DNeasy PowerSoil® Pro Kit, Qiagen, Hilden, Germany; Cat. No. 47014). Cell walls of bacteria were disrupted, and cells were lysed using a FastPrep FP120 cell homogenizer (Thermo Savant, USA) (20 seconds, 4°C, repeated 3 times with 2-minute intervals). DNA was then extracted using the DNeasy PowerSoil® Pro Kit (Qiagen, Hilden, Germany; Cat. No. 47014). The samples were submitted to the Integrated Microbiome Resource (Dalhousie University) for amplification and sequencing of V4-V5 regions of bacterial 16S rDNA and the V6-V8 regions of archaeal 16S rDNA. Data analysis followed Comeau et al. (2017), and results were visualized using QIIME2 (https://view.qiime2.org/).

#### Bacterial genome reassembly and annotation

Given Flye’s strength in assembling circular genomes, its assembly described above was used for endosymbiont genome analysis. Contigs corresponding to the *Centrionella anaeramoebae* genome were initially identified by performing a BLASTN search of the Flye assembly contigs against the NCBI nr database, considering features such as circularity and sequencing coverage similar to the host genome. To obtain a more accurate genome, Illumina data were aligned to the initially identified *Centrionella anaeramoebae* contigs using Bowtie2. These aligned reads, combined with the original Nanopore long reads, were then used for hybrid reassembly with Unicycler v.0.4.8, yielding the final circular genome of 1,521,317 bp.

The *Centrionella anaeramoebae* genome was gene called and annotated using NCBI Prokaryotic Genome Annotation Pipeline (2025-05-06.build7983) ^56^. The genome was annotated with 1249 genes (1198 CDSs, 44 tRNA, 3 rRNA, 4 ncRNAs) and 18 pseudogenes.

Three contigs belonging to *Desulfobacter* sp. LANTAAN genome was identified using similarity to *Desulfobacter* sp. BMAN and BMAN2. The long-read contigs were used to recruit short-reads and combined with nanopore reads in a hybrid reassembly using Unicycler as described for the endosymbiont genome. This yielded a circular genome of 7,469,478 bp. The genome was predicted and annotated using Bakta v1.11.4, yielding 6,352 CDSs, 70 tRNA, 18 rRNA, 4 ncRNA, and tmRNA^57,58^.

Anti-phage defense systems and toxin-antitoxin systems were predicted via the TADB3.0 server ^59^, PADLOC-DB v2.0.0 web server ^60^, and DefenseFinder v. 2.0.1 ^61,62^. Effector predictions were performed on the *C. anaeramoebae* PGAP predicted genome using the Searching algorithm for Type IV effector proteins (S4TE) 2.0 webserver using default settings (2025-10-12) ^25^.

### Bacterial expression analyses

The expression of genes in the bacterial genomes was performed using the salmon quant function in Salmon 1.11.4 ^63^ using an index prepared from the predicted transcripts. Both bacteria mapped relatively few reads (*C. anaeramoebae* – 24,669 fragments, *Desulfobacter* sp. LANTAAN – 27,541 fragments) as expected from a polyA-selected RNAseq library that would heavily deplete bacterial mRNAs. Given the low number of mapped fragments, the bacterial gene expression data needs to be interpreted with caution.

### Genome classification via GTDB-Tk

The classification of the *C. anaeramoebae* and *Desulfobacter* sp. LANTAAN used bac120 GTDB-Tk v2.6.1 using the full database release 226 downloaded on 2026-02-25, running on the Galaxy Europe servers. The *C. anaeramoebae* genome exhibited taxonomic novelty using as determined by a RED score of 0.89114. The genome was classified to _Bacteria;p Pseudomonadota;c Gammaproteobacteria;o UBA12402;f UBA12402;g JAF GHQ01;s. The *Desulfobacter* sp. LANTAAN taxonomic classification was defined by topology as d Bacteria;p Desulfobacterota;c Desulfobacteria;o Desulfobacterales;f Desulfobacterace ae;g Desulfobacter;s and had a RED value of 0.98147.

### Gene family and protein clustering

Gene family clustering performed using the OrthoVenn3 webserver ^64^ to cluster gene families in *C. anaeramoebae* and *Candidatus* Carsonella ruddii. The *C. anaeramoebae* proteome was clustered using CLANS v.29.05.2012 ^65^ at an e-value limit of 9.2E-5 to reveal additional protein families and to determine if there are shared domains. CLANS clusters were manually examined, and 15 clusters were annotated and visualized.

### Protein domain predictions in gene families

Domain annotations was performed of proteins families (n≥3) with InterProScan 5.77-108.0 against Pfam, NCBIfam, CDD, HAMAP, ProSite Profiles, ProSite Patterns, and SMART databases. Where multiple databases annotated overlapping regions, the highest-priority hit (Pfam > NCBIfam > CDD > SMART > ProSite) was retained after greedy non-overlapping selection. Of 263 proteins across 31 gene clusters (n≥3, 203 non-mobile and 60 mobile-element proteins are depicted; 66 of the non-mobile proteins (spanning 21 clusters) carry an S4TE prediction.

### Phylogenomic Analysis

Totally, 251 annotated protein sequences of *A. pumila* were separately combined with their orthologous protein sets from representative eukaryotic species and Metamonada protists. All data sets were aligned separately using the MAFFT algorithm ^66^, and ambiguous sites were removed with trimAl v.1.4 under default parameters ^67^. The alignment results of all protein sequences were then concatenated for phylogenetic tree construction. The Maximum Likelihood (ML) trees were built using IQ-TREE v.1.5.5, with the LG+C60+F+G4 model, and 100 replicates ^68^.

Following the phylogenetic analysis framework from ^11^, 109 protein sequences from the endosymbiont were separately combined with their corresponding orthologous protein sets, and then aligned and concatenated for phylogenetic analysis. The ML tree was constructed as described above. The Bayesian Inference (BI) trees were constructed using PhyloBayes v.4.1 under the CAT+GTR model, with 1,000,000 generations of Markov chain Monte Carlo sampling ^69^. Finally, the topology of the phylogenetic tree was visualized using FigTree v.1.4.4.

The phylogenomic analyses of *Desulfobacte*r sp. LANTAAN used the dataset and strategy described for phylogenomic reconstruction of *Anaeramoeba flamelloides* and *A. ignava Desulfobacter* sp. symbionts ^8^. The gene markers for the phylogenomic analysis of the Bac109 were identified with phyloSkeleton v1.160 ^70^. The UPF0081 marker was removed since it was only present in the out-group genome but not in any of the in-group genomes. The 108 proteins were aligned by MAFFT-linsi v7.458 ^66^ and trimmed by BMGE v1.12 ^71^, using the BLOSUM30 matrix and stationary-based character trimming. The final dataset consisted of 22,928 sites. The final dataset consisted of 36 genome-derived sequences: *Desulfovibrio desulfuricans* ND132 (1 genome), *Desulfobacteraceae* (23 genomes), *Desulfobacter* (8 genomes), and the *Anaeramoeba* symbionts or associated organisms (4 genomes). A maximum-likelihood tree was inferred using IQ-TREE v2.0.3 ^68^ using the LG + C20 + F + Gamma mixture model ^72^, and branch confidence was calculated by 1,000 ultra-fast bootstrap replicates and Shimodaira-Hasegawa approximate likelihood-ratio test.

### Fluorescence In Situ Hybridization (FISH)

#### Design of Specific FISH Probes for Endosymbiont

The endosymbiont genome was submitted to the RNAmmer 1.2 Server for 16S rDNA prediction (Lagesen et al., 2007), and primers were designed. PCR amplification of 16S rDNA was performed using Thermo Scientific Phusion Hot Start II High-Fidelity DNA Polymerase (Thermo Fisher Scientific, USA, Cat. No. F549L), and the 16S rDNA was sequenced by GENEWIZ.

The 16S rDNA sequence was aligned with those of 47 other representative bacterial species (downloaded from the Silva High Quality Ribosomal RNA Database, https://www.arb-silva.de/browser/) using MAFFT ^66^. The alignment was submitted to DECIPHER for probe design ^73^, and probe specificity was verified using the RDP databases (https://rdp.cme.msu.edu/index.jsp).

#### FISH Experiment

Firstly, the amoeba cells were collected by removing supernatant, tapping the tube, and washing with ASW. Then, the remaining liquid (∼200 µL) was used to resuspend and was quickly added to a 6-well printed slide (”PTFE” Printed Slides (ER-264W), Electron Microscopy Sciences, USA, Cat. No. 63423-08) at 100 µL per well and allowed to adhere for 5 minutes at room temperature. Cells were washed twice with ASW, then fixed with an appropriate volume of 16% methanol-free formaldehyde solution (FA) (Thermo Fisher Scientific, USA, Cat. No. 28906) to achieve a final concentration of approximately 4% for 10 minutes at room temperature. Slides were washed twice with distilled water (1.5 minutes each), air-dried, and examined under a microscope. Dehydration was performed with 95% ethanol for 5 minutes, followed by air-drying and microscopic examination. FISH hybridization buffer preparation and experimental procedures followed ^74^. Finally, ProLong™ Diamond Antifade Mountant with DAPI (Thermo Fisher Scientific, Cat. No. P36962) was added to each well, and coverslips (Fisher brand® Microscope Cover Glass, Thermo Fisher Scientific, USA, Cat. No. 12544E) were applied for sealing.

Slides were incubated in the dark at room temperature for 12 hours. Microscopic images were captured using either a Zeiss Axiocam 506 mono Imager Z2 fluorescence microscope or a Leica SP8 confocal microscope (Cellular & Molecular Digital Imaging Facility, Dalhousie University).

### Immunofluorescence staining

Processing of *A. pumila* cells for immunofluorescence staining, slide preparation, and dehydration was performed as described in the FISH experiment section above. If FISH and IF were combined, the FISH protocol was followed until before the mounting, and the slides were post-fixed in 4% FA for 10 min, washed twice with distilled water (1.5 minutes each), followed by quenching using 50 mM NH4Cl to the remaining aldehyde fixative.

Subsequently, each well was washed twice with PBS buffer. Cells were incubated with Antibody Blocking Buffer (ABB Buffer: 1% AURION BSA-c™ 10% (Electron Microscopy Sciences, USA, Cat. No. 25557) + 0.1% Triton® X-100 (Sigma, USA, Cat. No. T9284) + PBS) for 1 hour at room temperature. Two heterologous antibodies, α-tubulin TAT1 and *Neocallimastix* Chaperonin 60 (NeoCpn60) ^75^, were incubated for 1 hour at room temperature. TAT1 was diluted 1:200 in ABB Buffer, and the NeoCpn60 was diluted 1:1000 in ABB Buffer. After incubation, cells were washed eight times with PBS buffer. Secondary antibody solutions were applied as appropriate and incubated in the dark at room temperature for 1 hour. For TAT1, the secondary antibody was Alexa Fluor® 594 goat anti-mouse IgG (H+L) (2 mg/mL, Thermo Fisher Scientific, Cat. No. A-11032) diluted 1:200 in ABB Buffer. For NeoCpn60, the secondary antibodies were either Alexa Fluor™ Plus 488 goat anti-rabbit IgG (H+L) (2 mg/mL, Thermo Fisher Scientific, Cat. No. A-32731) or Alexa Fluor™ Plus 647 goat anti-rabbit IgG (H+L) (2 mg/mL, Thermo Fisher Scientific, Cat. No. A-32733), both diluted 1:200 in ABB Buffer. After eight washes with PBS buffer, slides were mounted and imaged as above.

### Focused-ion-beam scanning electron microscopy (FIB-SEM) and segmentation

Cells were adhered to the surface of a gridded MatTek dish and fixed with 2.5% Glutaraldehyde (TAAB) in 0.1M PHEM-buffer. All samples were processed using Pelco Biowave Pro+ microwave tissue processor (Ted Pella, Redding, CA) according to ^76^ with minor modifications: no calcium was used during fixation, and the contrasting steps with lead aspartate were omitted to reduce the risk of overstaining. The sample was embedded in Durcupan resin (Sigma (Merck Life Science, Solna, Sweden). The embedded block was separated from the glass using liquid nitrogen and glued to an SEM-stub with epoxy and silver glue. The sample was further coated with 5nm Pt to reduce charging. Volumes were acquired using a Scios dualbeam (FEI, Eindhoven, The Netherlands) with the electron beam operating at 2 kV/0.2 nA detected with the T1 In-lens detector. To automate the acquisition, we used the Auto Slice and View 4 software provided with the microscope. A 700nm protective layer of platinum was deposited on the selected area before milling. The volume was further registered and processed by the ImageJ plugins Linear alignment by SIFT and Multistackreg. After registration, the volumes were converted to mrc-files, and the header was modified to recover the pixel size that got lost during conversion. The final image consisted of 1580 slices with a voxel size of (0.00562 µm × 0.00562 µm × 0.06 µm).

### Segmentation

The *A. pumila* cell was manually segmented in Microscopy Image Browser v2.91 beta 45 using Segment Anything Model 2 (SAM 2). The segmentations included the nucleus, four symbiont clusters, microtubule-organizing centers, and hydrogenosomes. The segmentations were rendered in Dragonfly v.2022.2.0 Build 1399.

## Data availability

Sequencing reads and the annotated genome of *Anaeramoeba pumila* LANTAAN, *Centrionella anaeramoebae*, and *Desulfobacter* sp. LANTAAN were deposited to NCBI under the BioProject numbers PRJNAXXXXXXX, PRJNAXXXXXXX, and PRJNAXXXXXXX, respectively. The RNAseq reads, genomic DNA Illumina reads, and nanopore reads are available from SRA under numbers SRRXXXXXXXX, SRRXXXXXXXX, and SRRXXXXXXXX, respectively. FIB-SEM aligned mrc-files and segmentation models in MODEL format are available from Figshare (10.6084/m9.figshare.31866142).

## Acknowledgements

The authors would like to acknowledge Umeå Centre for Electron Microscopy (UCEM) for technical assistance and access to electron microscopy. Service/Support/Collaboration was provided/supported by SciLifeLab Integrated Microscopy Technology Unit at Umeå University.

## Author contributions statement

Conceptualization, A.J.R., and J.J-H.; Investigation, T.Z., and M.P., J.J-H.; Formal analysis, T.Z., D.S.L., S.W. and J.J-H.; Resources, I.Č., A.J.R. and J.J-H.; Writing – Original Draft, J.J-H.; Writing – Review & Editing, T.Z., M.P., D.S.L., S.W., I.Č., A.J.R. and J.J-H.; Funding Acquisition, A.J.R. and J.J-H.

## Figures and Supplementary material

**Figure S1:**
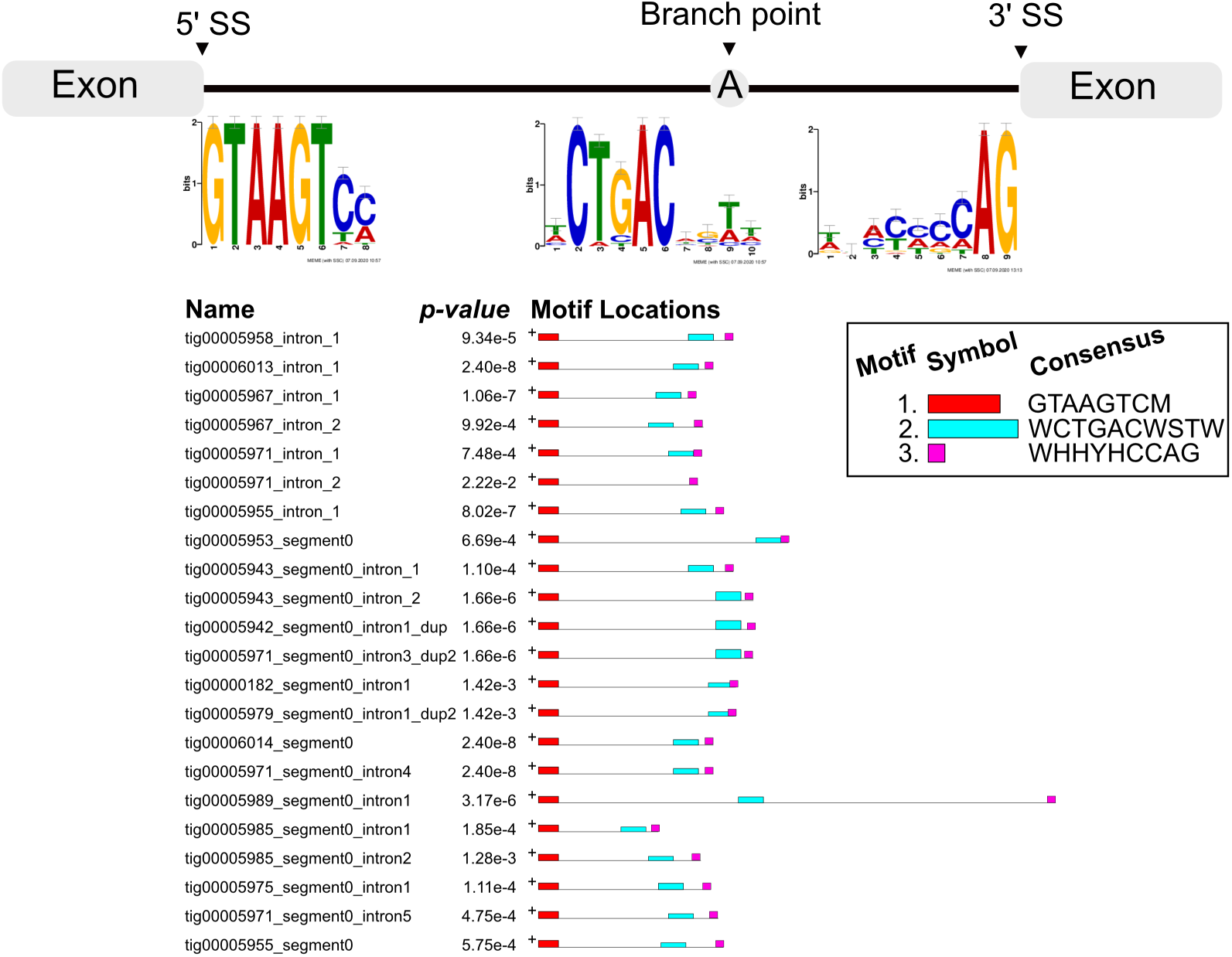
*A. pumila* has few and structured splicing motifs. List showing 22 introns in 19 genes seen in the *A. pumila* genome. Three genes have two introns each, and a further three introns are 100% redundant (2 copies, 3 copies, 3 copies), examples from gene duplicates. This leaves 17 unique splicing examples with an average size of 79.9 bp, ranging between 48 bp and 207 bp. The average GC content is 45%, which is slightly below the genome average of 52-53%. All introns were supported by spliced RNA-seq reads and yield functional proteins when spliced *in silico*. The 5’ splice site shows a highly conserved profile (5’|GTTAGT). The 3’ end showed no clear signs of conservation besides the AG dinucleotide. There is conservation around the predicted branch-point A-nucleotide (CT(G/c)**A**C). The conserved patterns were calculated using MEME^7^. Abbreviations: 5’SS, 5’ splice site; 3’SS, 3’ splice site.

**Figure S2:**
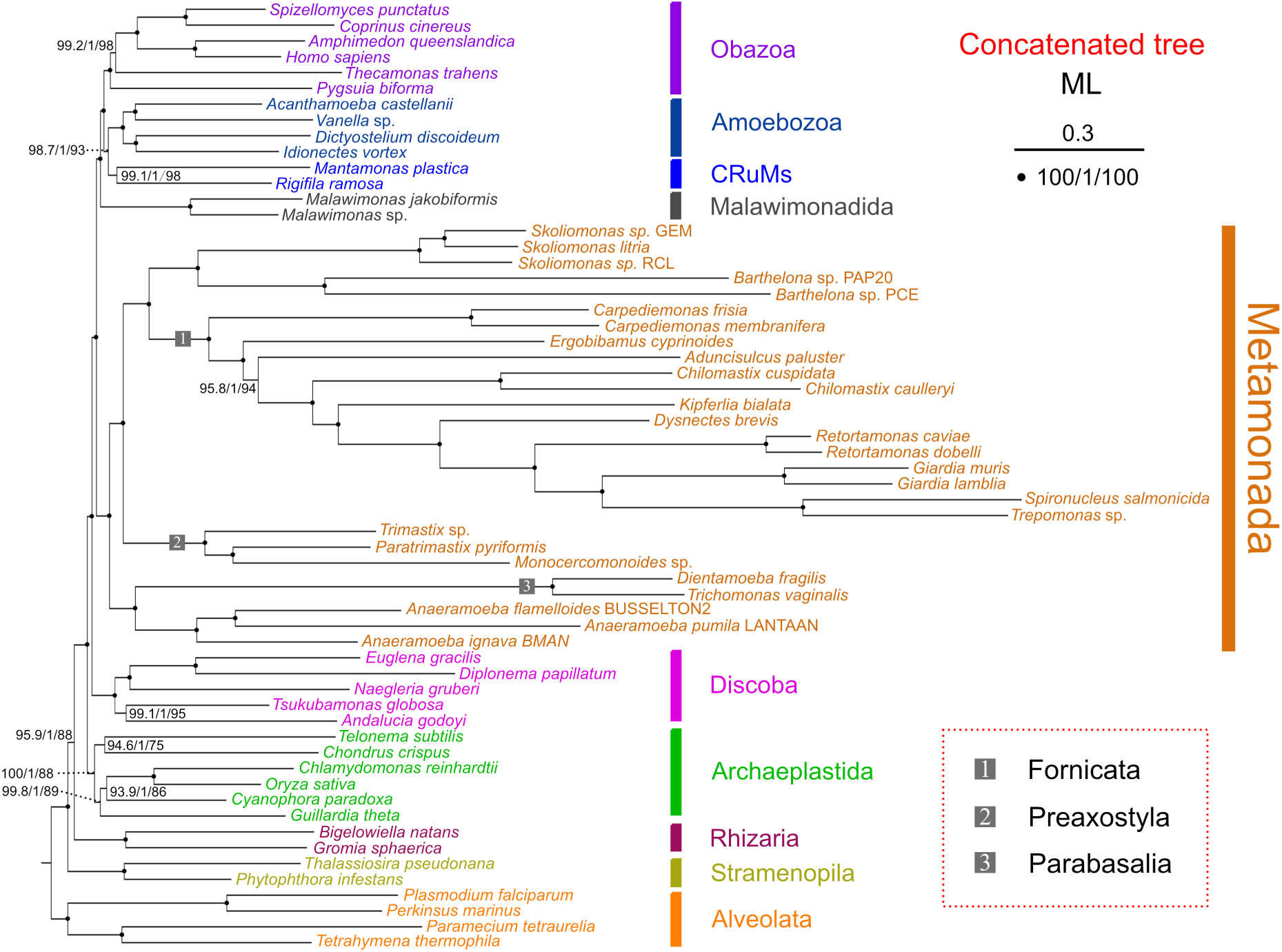
The maximum likelihood tree (ML) focusing on the Metamonada, generated from 252 protein sequences with a total of 65,588 amino acid sites. The ML tree was constructed using IQ-TREE with the model LG+C60+F+G4. Values at nodes represent SH-aLRT, aBayes, and standard confidence values, respectively. The solid circles indicate full confidence values. The scale bar indicates a substitution rate of 30%.

**Figure S3:**
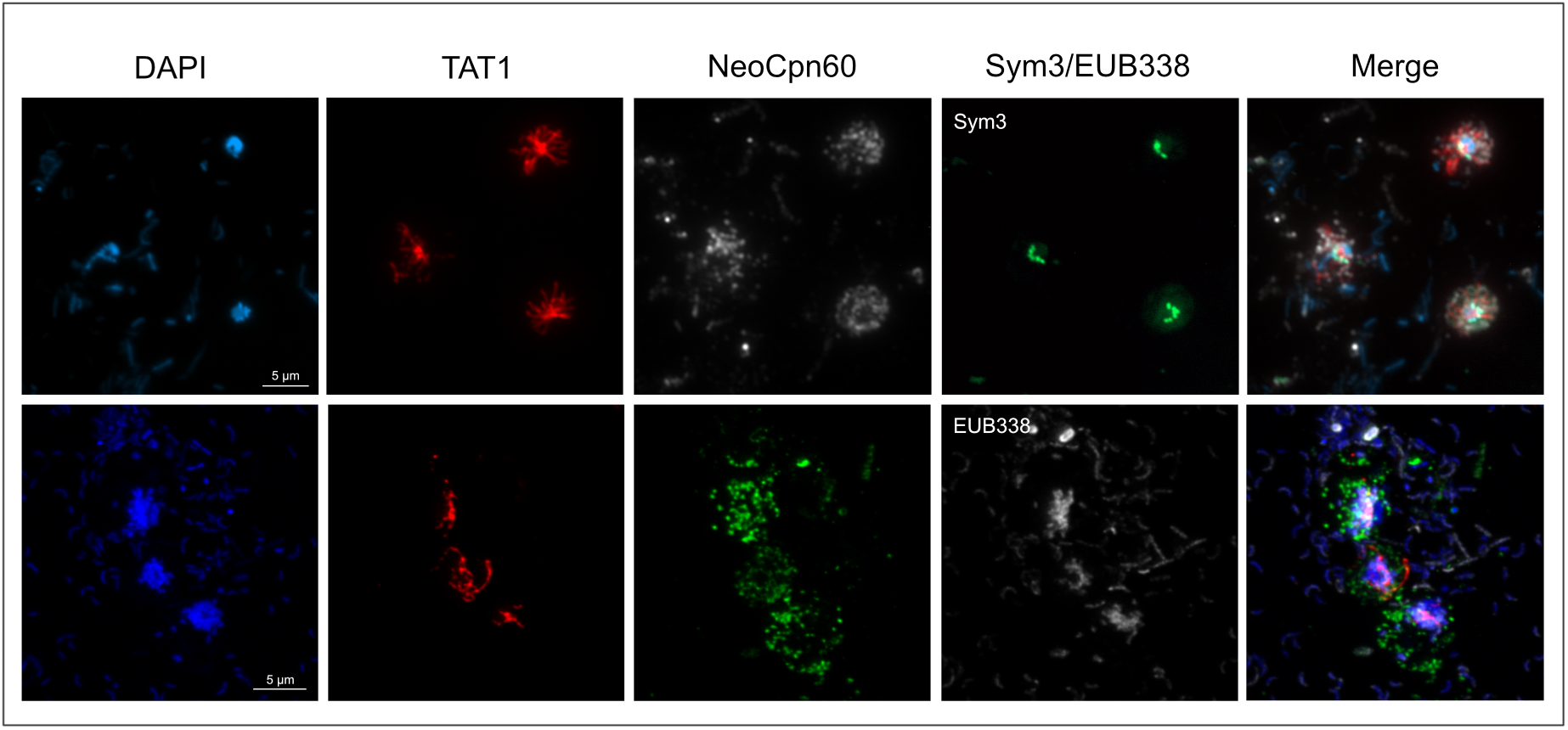
Immunofluorescence staining of mitochondrion-related organelles (MROs) in *A. pumila*. First column (blue channel): Host nuclei and symbiont DNA were stained using DAPI. Second column (red channel): Immunofluorescence staining of alpha-tubulin (TAT1 antibody, 1:200), labels the acentriolar centrosome and radiating microtubule. Third column (white/green channels): Putative MROs were stained using a heterologous *Neocallimastix* Cpn60 (NeoCpn60) antibody^8^. The signal shows MRO-resident chaperonin Cpn60 shows a punctuate distribution. Different secondary antibodies were used to enable the simultaneous detection of Sym3 and EUB338. Fourth column (Green/White channel): The *C. anaeramoebae*-specific Sym3 (green) and universal bacterial probe EUB338 (white) label the symbiont and bacteria, respectively. Fifth column: A merged image of columns 1-4. Scale bar 5 µm.

**Figure S4:**
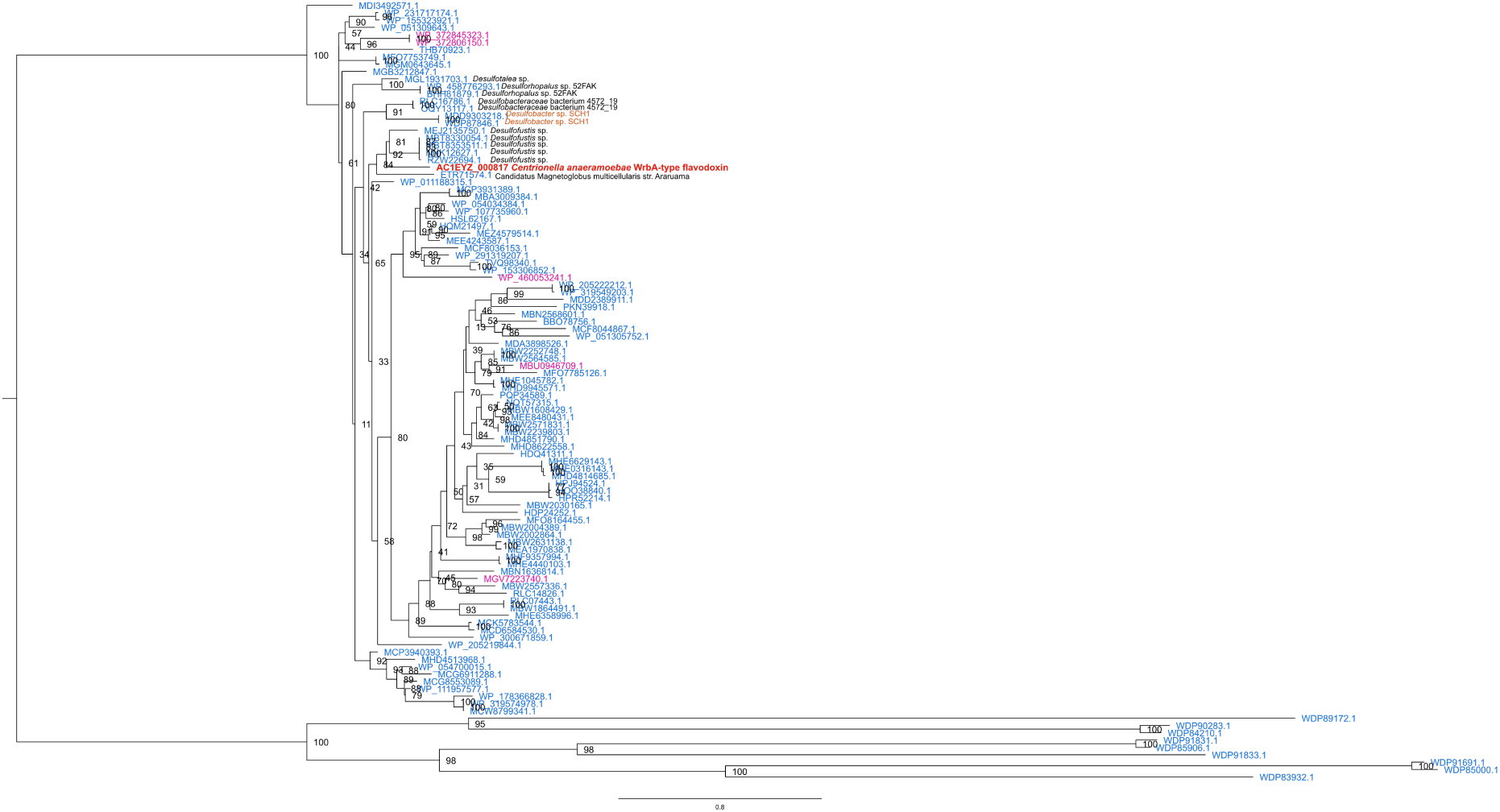
Lateral gene transfer of WrbA-like flavodoxins from Desulfobulbales. Phylogenetic analyses of WrbA-like flavodoxin in the *Centrionella anaeramoebae* genome. The ML tree (106 sequences with 170 amino-acid sites (158 informative)) was constructed using IQ-TREE 3.1.0 with the model LG4X+I+R5 as selected by ModelFinder. Values at nodes represent ultrafast bootstraps (1000 replicates).

**Figure S5:**
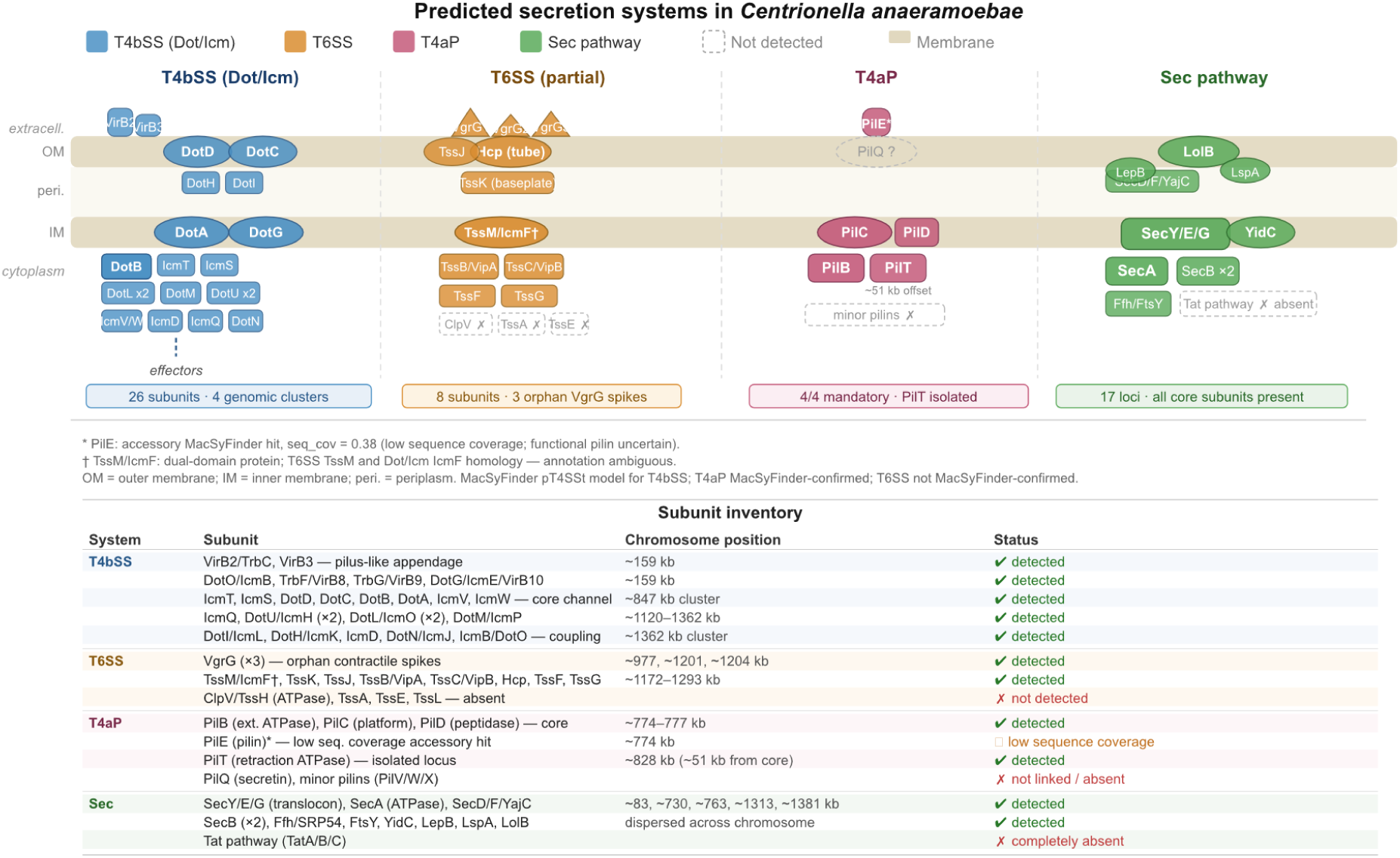
Secretion systems in *Centrionella anaeramoebae*. Schematic representation of four distinct protein secretion systems detected in *C. anaeramoebae* through comprehensive genomic analysis. Systems are organized by cellular location relative to the outer membrane (OM), periplasm, and inner membrane (IM), with detection confidence indicated for each system type. Shaded shapes represent detected subunits; dashed outlines indicate absent or unlinked components. **T4BSS Dot/Icm (Type IVb)**: Complete multiprotein complex spanning both membranes. Key components include VirB2/3 (surface), DotD/C (periplasmic), DotA/G (IM ATPases), and DotB (cytoplasmic motor). 26 subunits detected across 4 genomic clusters (159, 847, 1120, 1362 kb). **T6SS (Type VI, partial)**: Degenerate system with 8/13 core components. Present: VgrG×3 (orphan spikes), Hcp tube protein, TssJ-K-M baseplate, TssB/C sheath, TssF/G. Absent: ClpV ATPase, TssA/E/L assembly factors. Located at 977, 1172, 1287 kb. **T4aP (Type IVa pilus)**: All 4 mandatory motor components present: PilB (extension ATPase), PilC (alignment platform), PilD (peptidase), PilT (retraction ATPase at 828 kb). Core apparatus at 773–777 kb. PilE pilin has low sequence coverage; minor pilins absent. **Sec pathway (General secretion)**: Complete system with all 17 core loci: SecYEG translocon, SecA motor, SecDFYajC holotranslocon, YidC insertase, SRP machinery (Ffh/FtsY), signal peptidases (LepB/LspA), SecB chaperones, and LolB lipoprotein insertase. Tat pathway completely absent. Secretion system detection: MacSyFinder TXSScan with manual verification for divergent systems. Notes: PilE marked with asterisk due to low sequence coverage (0.38). TssM/IcmF annotation is ambiguous due to dual homology. Shaded shapes represent confirmed subunits; dashed outlines indicate absent or MacSyFinder-unlinked components. OM = outer membrane; IM = inner membrane.

**Figure S6:**
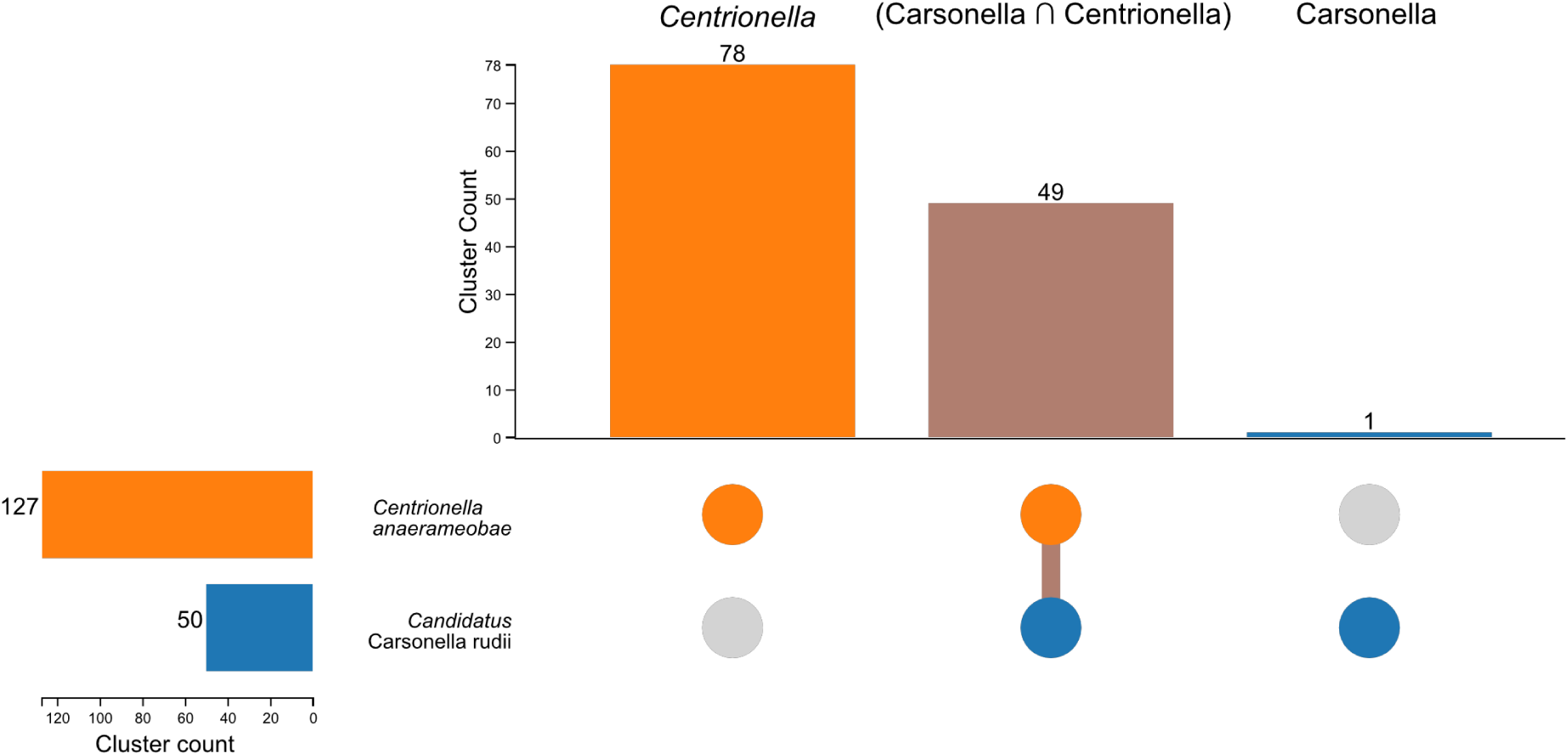
Gene family clustering of *Centrionella anaeramoebae* and *Candidatus* Carsonella ruddii genomes. OrthoVenn3 was used to cluster the proteomes of *C. anaeramoebae* and *Candidatus* Carsonella ruddii. **(A)** Upset-plot of the clustered gene families across the two genomes. The clustering revealed 78 gene families, a total of 156 proteins with more than 2 members or more.

**Figure S7:**
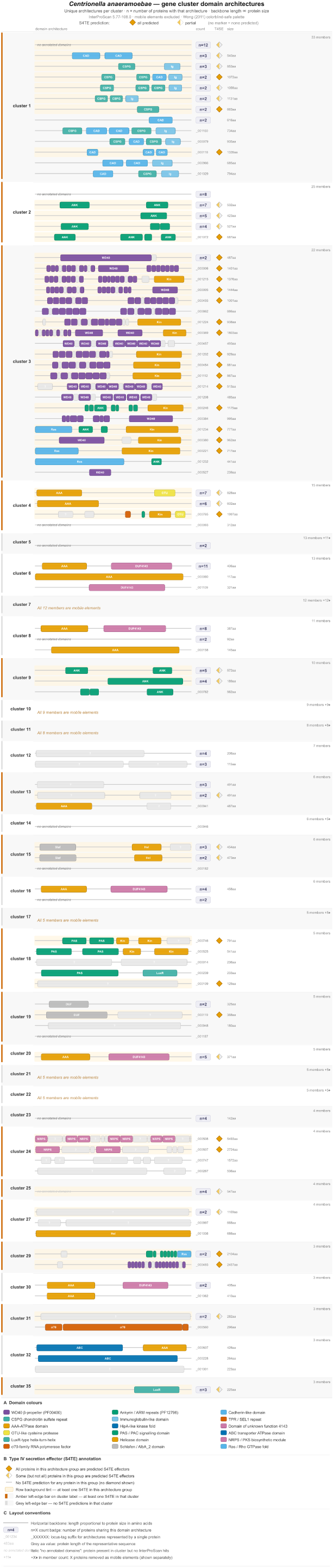
Domain architecture of *Centrionella anaeramoebae* gene cluster proteins with predicted type IV secretion effectors. Each row represents a unique domain architecture observed among proteins in a given gene cluster. Proteins sharing identical domain order and composition are collapsed into a single row, with the count of member proteins indicated by the n = value to the right of the backbone. Rows represented by a single protein show the locus tag suffix in grey. The horizontal backbone is proportional to protein length (amino acids; shown at the right margin). Proteins for which InterProScan returned no domain annotation are indicated by italic ‘no annotated domains’. Mobile element proteins (transposases and integrases) are excluded from this figure; their count per cluster is appended as +n♦ in the member tally. **Domain color scheme.** Colored boxes indicate annotated protein domains drawn to scale within each backbone. The following domains are shown: **WD40 β-propeller** (PF00400; purple). **Ankyrin / ARM repeats** (PF12796; bluish green). **Cadherin-like domain** (sky blue). **CSPG chondroitin sulfate repeat** (teal). **Immunoglobulin-like domain** (light blue). **TPR / SEL1 repeat** (vermillion). **AAA-ATPase domain** (amber; shared color with Helicase). **HipA-like kinase fold** (blue; shared color with ABC transporter). **Domain of unknown function 4143 (DUF4143)** (reddish purple; shared color with NRPS/PKS). **OTU-like cysteine protease** (yellow). **PAS / PAC signalling domain** (bluish green; shared color with Ankyrin). **ABC transporter ATPase domain** (blue; shared color with HipA). **LuxR-type helix-turn-helix** (teal; shared color with CSPG). **Helicase domain** (amber; shared color with AAA-ATPase). **NRPS / PKS biosynthetic module** (reddish purple; shared color with DUF4143). **σ70-family RNA polymerase sigma factor** (vermillion; shared color with TPR). S4TE effector predictions are encoded at three levels of resolution. A filled amber diamond (◆) to the right of the count badge indicates that **all** proteins sharing that architecture are predicted S4TE effectors. A half-filled diamond (◆, amber/white) indicates that **some but not all** proteins in the group carry a S4TE prediction; no diamond is shown when none are predicted. Rows containing at least one S4TE protein are highlighted with a pale amber background. Cluster labels bearing an amber left-edge bar contain at least one S4TE-predicted protein among their non-mobile members; a grey bar indicates no S4TE predictions for that cluster.

**Figure S8:**
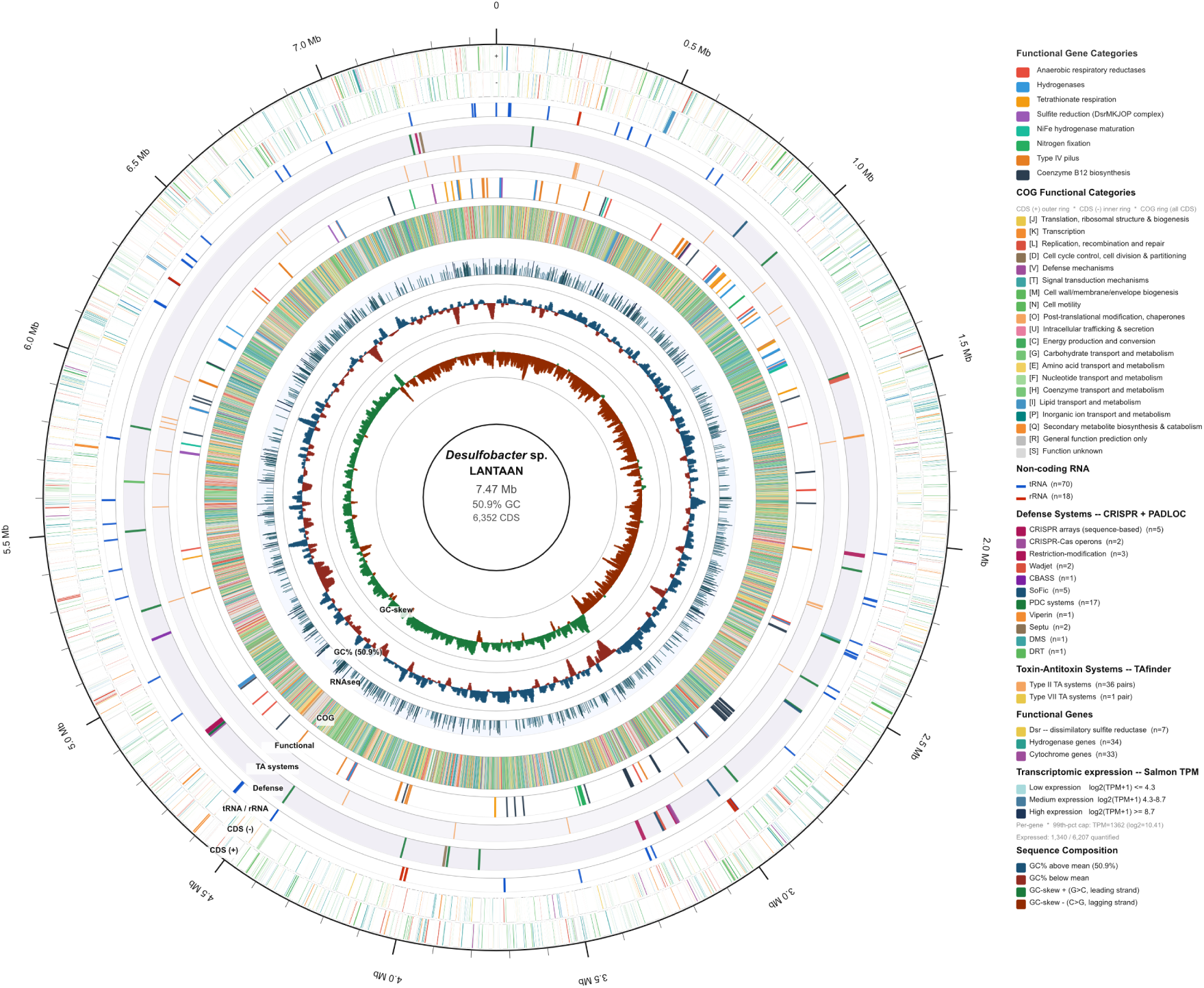
Genome atlas of *Desulfobacter* sp. LANTAAN. Circular genome map of the *Desulfobacter* sp. LANTAAN chromosome (7,469,478 bp; 50.9% GC). Eleven concentric tracks are arranged from outer to inner and are described in detail below. All tracks are drawn at exact genomic coordinates; no positional smoothing or gene-length normalisation was applied. The annotation comprises 6,352 protein-coding sequences, 70 tRNA genes, 18 rRNA genes (multiple 16S, 23S, and 5S copies), and 5 CRISPR arrays (sequence-based repeats). **Track 1. Scale and tick marks.** The outermost ring shows the chromosomal coordinate scale. Major tick marks are placed every 500 kb with Mb labels rendered as tangent-rotated text (font-weight 500, size 11 pt, #1A1A1A). Minor tick marks are placed every 100 kb. Coordinates are numbered clockwise from the origin (position 0 at the top) and correspond directly to the Bakta annotation coordinates. **Track 2 and 3.** Protein-coding sequences, forward (+) strand and, reverse strand, respectively. **Track 4.** Non-coding RNA genes. This narrow ring marks the positions of the 70 tRNA genes (blue, #1155CC) and 18 rRNA genes (dark red, #CC2200). Source: Bakta v1.11.4 GenBank tRNA and rRNA features. **Track 5. Defense systems — PADLOC.** This track displays all 35 defense systems predicted by PADLOC v.2.0.0^1^, plus the 5 CRISPR arrays identified from sequence. Each system type is assigned a distinct color. **Track 6. Toxin-antitoxin systems — TAfinder 2.0.** This track displays the 37 toxin-antitoxin (TA) pairs (74 total genes) predicted by TAfinder 2.0^2^. Type II TA systems (n = 36 pairs). Type VII TA systems (n = 1 pair). Toxin and antitoxin components of each pair are co-positioned and share the same color. **Track 7. Functional gene highlights.** This track displays a curated catalogue of 230 functional genes across eight metabolic and cellular categories. Anaerobic respiratory reductases (58 genes), Hydrogenases (35 genes), Tetrathionate respiration (14 genes), Sulfite reduction-associated complex DsrMKJOP(12 genes), NiFe hydrogenase maturation (7 genes), Nitrogen fixation (13 genes), Type IV pilus (34 genes), Coenzyme B12 biosynthesis (57 genes). The categories were annotated using the Rapid Annotations using Subsystems Technology (RAST) server^9^. **Track 8. COG functional annotation ring**. A narrow ring repeats the COG color code for all 6,352 CDS (both strands combined), providing a rapid overview of functional composition around the chromosome. Colors are identical to those used in **Tracks 2** and **3**. COG categories were assigned by keyword matching against product annotations; genes whose products could not be assigned to an information-processing, metabolic, or cellular-process category were classified as R (general function prediction only) or S (function unknown). Where eggNOG-mapper assigned multiple COG letters to a gene (e.g., “NT”), the first letter was used as the primary assignment. Colors are identical to those used in **Tracks 2** and **3** (see COG color key, below). **Track 9. Transcriptomic expression — Salmon TPM**. This histogram track displays per-gene transcript abundance estimated by Salmon quantification^4^. Values are expressed as transcripts per million (TPM) and plotted on a log₂(TPM + 1) scale. To prevent extreme outliers from compressing the color gradient, values are capped at the 99th-percentile TPM (921; log₂ = 9.85). Of the 6,207 genes quantified, 1,340 had TPM > 0 (expressed) and 4,867 had TPM = 0 (not detected). The most highly expressed gene is 2294.16_05277 (PEP-CTERM sorting domain protein; TPM = 103,795); Bar height encodes the normalised expression level (0–1 scale). Bar color transitions continuously from pale teal through steel blue to deep navy as expression increases. **Track 10. GC content** GC percentage was calculated in 15-kb sliding windows with a 3-kb step (2,485 windows total), producing a mean GC of 50.9% (range 33.1–60.4%). The histogram baseline is set at the genome-wide mean; windows above the mean are drawn in dark blue (#1A5276) and windows below the mean are drawn in dark red (#922B21), making regional compositional biases immediately apparent. A dashed blue baseline marks the mean GC level. **Track 11. GC-skew.** GC-skew, defined as (G − C) / (G + C), was calculated over the same 15-kb/3-kb windows as Track 10 (range −0.146 to +0.180). The baseline is set at zero; windows with G surplus (positive skew) are drawn in dark green (#1A7A3C) and windows with C surplus (negative skew) are drawn in dark orange (#922B00). The sign transitions of GC-skew correspond to the putative replication terminus (negative-to-positive, Δ at ∼3.7 Mb) and origin of replication (positive-to-negative, Δ at ∼7.1/0.0 Mb), consistent with a single bidirectional origin in this organism.

